# Vascular control of the CO_2_/H^+^ dependent drive to breathe

**DOI:** 10.1101/2020.08.19.257386

**Authors:** CM Cleary, TS Moreira, AC Takakura, MT Nelson, TA Longden, DK Mulkey

## Abstract

Respiratory chemoreceptors regulate breathing in response to changes in tissue CO_2_/H^+^. Blood flow is a fundamental determinant of tissue CO_2_/H^+^, yet little is known regarding how regulation of vascular tone in chemoreceptor regions contributes to respiratory behavior. Previously, we showed in rat that CO_2_/H^+^-vasoconstriction in the retrotrapezoid nucleus (RTN) supports chemoreception by a purinergic-dependent mechanism (Hawkins et al. 2017). Here, we show in mice that CO_2_/H^+^ dilates arterioles in other chemoreceptor regions, thus demonstrating CO_2_/H^+^ vascular reactivity in the RTN is unique. We also identify P2Y_2_ receptors in RTN smooth muscle cells as the substrate responsible for this response. Specifically, pharmacological blockade or genetic deletion of P2Y_2_ from smooth muscle cells blunted the ventilatory response to CO_2_, and re-expression of P2Y_2_ receptors only in RTN smooth muscle cells fully rescued the CO_2_/H^+^ chemoreflex. These results identify P2Y_2_ receptors in RTN smooth muscle cells as requisite determinants of respiratory chemoreception.

**Significance Statement:** Disruption of vascular control as occurs in cardiovascular disease leads to compromised chemoreceptor function and unstable breathing. Despite this, virtually nothing is known regarding how regulation of vascular tone in chemoreceptor regions contributes to respiratory behavior. Here, we identify P2Y_2_ receptors in RTN vascular smooth muscle cells as a novel vascular element of respiratory chemoreception. Identification of this mechanism may facilitate development of treatments for breathing problems including those associated with cardiovascular disease.

## Introduction

Blood flow in the brain is tightly coupled to neural activity. Typically, an increase in neural activity triggers vasodilation to increase delivery of nutrients and clear metabolic waste (6, 21). Blood flow may also directly regulate neural activity and contribute to behavior; perhaps the most compelling evidence supporting this involves vasoregulation by CO_2_/H^+^. In most cases, CO_2_/H^+^ function as potent vasodilators to provide support for metabolic activity and to maintain brain pH within the narrow range that is conducive to life (1). However, CO_2_/H^+^ has the opposite effect on vascular tone in a brainstem region known as the retrotrapezoid nucleus (RTN) (18, 40), where CO_2_/H^+^ is not just metabolic waste to be removed but also functions as a primary stimulus for breathing (17). Neurons (17, 22) and astrocytes (14) in this region are specialized to regulate breathing in response to changes in tissue CO_2_/H^+^ (i.e., function as respiratory chemoreceptors) and so vasoconstriction may augment this function by preventing CO_2_/H^+^ washout and maintaining the stimulus to respiratory chemoreceptors. However, it is not known whether CO_2_/H^+^ induced vasoconstriction is unique to the RTN or also contributes to chemotransduction in other chemosensitive brain regions. Also, mechanisms underlying this specialized CO_2_/H^+^ regulation of vascular tone in the RTN are largely unknown. Previous work showed that RTN chemoreception involves CO_2_/H^+^-evoked ATP-purinergic signaling most likely from astrocytes (14, 39); however, downstream cellular and molecular targets of this signal are not known.

Here, we show that CO_2_/H^+^ vascular reactivity in the RTN is unique compared to other chemoreceptor regions. Our evidence also suggests that CO_2_/H^+^ constriction in the RTN enhances chemoreceptor activity and respiratory output, while simultaneous dilation in other chemoreceptor regions may serve to limit chemoreceptor activity and stabilize breathing. We also show that endothelial cells and vascular smooth muscle cells in the RTN have a unique purinergic (P2) receptor expression profile compared to other levels of the respiratory circuit that is consistent with P2Y_2_ dependent vasoconstriction. Further, at the functional level, we show pharmacologically and genetically using P2Y_2_ conditional knockout (P2Y_2_ cKO) mice and RTN smooth muscle-specific P2Y_2_ re-expression that P2Y_2_ receptors on vascular smooth muscle cells in the RTN are required for CO_2_/H^+^ dependent modulation of breathing.

## Results

### P2Y_2_ receptors are differentially expressed by vascular smooth muscle cells in the RTN

Purinergic signaling is required for CO_2_/H^+^-induced constriction of RTN arterioles(18). Therefore, as a first step towards identifying the cellular and molecular basis of this response, we determined the repertoire of P2 receptors expressed by endothelial and smooth muscle cells associated with arterioles in the RTN. We also characterize the P2 receptor expression profile of vascular cells in the caudal aspect of the nucleus of the solitary tract (cNTS) and raphe obscurus (ROb), two putative chemoreceptor regions where P2 receptor signaling does not contribute to respiratory output (35). For this experiment, we used Cre-dependent smooth muscle (smMHC^Cre/eGFP^) and endothelial cell (Tie2^Cre^::TdTomato) reporter lines and performed fluorescence activated cell sorting to obtain enriched populations of endothelial or smooth muscle cells. Control tissue was obtained at the same level of brainstem but outside the region of interest. Each population of cells was pooled from 3-8 adult mice per region for subsequent quantitative RT-PCR for each of the fourteen murine P2 receptor subtypes. The type and relative abundance of P2 receptors expressed by each cell type was similar to the cortex (38) and relatively consistent between brainstem regions. For example, endothelial cells from each region express ionotropic P2X_4_ and P2X_7_ and metabotropic P2Y_1,2,6,12-14_ receptors, albeit at varying levels relative to control (**Fig. 1A**). Also, P2X_1,4_ and P2Y_2,6, and 14_ receptor subtypes were detected in smooth muscle cells from each region and most were under-expressed relative to control (**Fig. 1B**). The P2 transcript profile of vascular cells in the RTN was fairly similar to the ROb (Figs. 1A-B) but differed from the cNTS in several ways including lower expression of P2Y_6_ in endothelial cells (**Fig. 1A**) and higher expression of P2X_1_, P2X_4_ and P2Y_14_ in smooth muscle cells (**Fig. 1B**). However, only P2Y_2_ in the RTN showed a distinct expression profile that was consistent across cell types with a role in vasoconstriction; P2Y_2_ transcript was detected at lower (−9.2 ± 0.3 log_2_ fold change, p=0.0013) and higher (6.8 ± 1.5 log_2_ fold change, p=0.0462) levels compared to control in RTN endothelial and vascular smooth muscle cells, respectively (**Figs. 1A-B**). This is interesting because in other brain regions activation of endothelial P2Y_2_ receptors favors nitric oxide-induced vasodilation (27), whereas activation of this same receptor in vascular smooth muscle cells mediates vasoconstriction (3, 23). These results identify P2Y_2_ receptors in vascular smooth muscle cells as potential substrates for CO_2_/H^+^-induced ATP-dependent vasoconstriction in the RTN.

**Figure 1.**
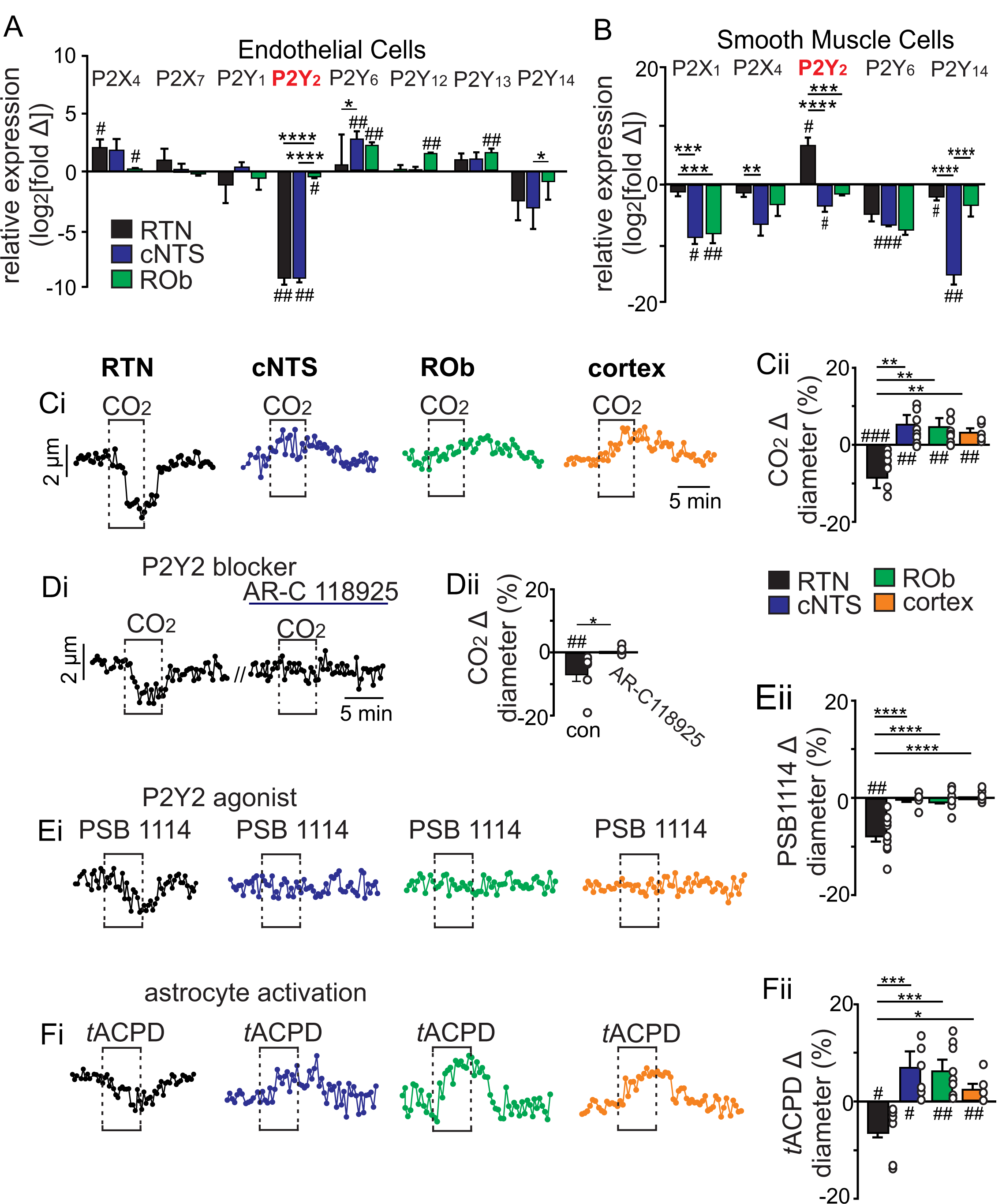
CO_2_/H^+^ differentially regulates arteriole tone in the RTN by a P2Y_2_ receptor dependent mechanism. **A-B**, tissue sections containing the cNTS, ROb, and RTN were prepared from an endothelial cell (Tie2^Cre^::TdTomato) and smooth muscle cell reporter mice (smMHC^Cre::eGFP^). Individual cells were dissociated and sorted to isolate enriched cell populations from each region (100-300 cells/region). Control cells for each region were prepared from slices with experimental regions removed. Fold change of each P2 receptor was determined for each region by normalizing to within group *Gapdh* expression as well as to control group receptor expression (**Supplemental Table 1**) and plotted as log_2_[fold change]; 0 on the y-axis indicates control group expression for each receptor and negative values reflect less than the control group and positive values reflect greater than control group. Of all P2 receptors detected in each region, only P2Y_2_ showed an expression pattern that was consistent with a role in vasoconstriction in the RTN; low expression in endothelial cells (**A**) and above baseline expression in smooth muscle cells (**B**) from this region. **C-F**, diameter of arterioles in the RTN, cNTS, ROb and somatosenstory cortex was monitored in brain slices from adult mice over time by fluorescent video microscopy. **C-F**, Diameter traces of individual arterioles in each region and corresponding summary bar graphs show that exposure to 15% CO_2_ (**Ci**), activation of P2Y_2_ receptors by bath application of PSB1114 (100 µM) (**Ei**), or activation of astrocytes by bath application of t-ACPD (50 μM) (**Fi**) caused vasoconstriction in the RTN and dilation in all other regions of interest. **Di-Dii**, The CO_2_/H^+^ response of RTN vessels was blocked by a selective P2Y_2_ receptor antagonist (AR-C118925; 10 μM) #, difference from baseline (paired t-test). *, differences in each condition (one-way ANOVA with Tukey’s multiple comparison test). One symbol = p < 0.05, two symbols = p < 0.01, three symbols = p < 0.001, four symbols = p < 0.0001.

### CO_2_H^+^ differentially effects vascular tone in the RTN by a P2Y_2_ dependent mechanism

To further test this possibility, we used the brain slice preparation, optimized for detecting increases or decreases in vascular tone, to characterize CO_2_/H^+^ vascular responses in each chemoreceptor region of interest as well in a non-respiratory region of the somatosensory cortex. For these experiments, we target arterioles based on previously described criteria (9, 29). We verified that exposure to CO_2_/H^+^ (15% CO_2_; pH 6.9) under control conditions decreases the diameter of RTN arterioles by -11.2 ± 4.5% (N=9 vessels), whereas this same stimulus consistently dilates arterioles in the cNTS (Δ +6.0 ± 2.2%; N=9 vessels), ROb (Δ +6.5 ± 2.6%; N=11 vessels) and somatosensory cortex (Δ +3.7 ± 0.9%; N=9 vessels) (F_3,34_=13.49; p <0.0001) (**Figs. 1Ci-Cii**). Consistent with a requisite role of P2Y_2_ receptors in this response, we found that CO_2_/H^+^-induced constriction of RTN arterioles was eliminated by blockade of P2Y_2_ receptors with AR-C118925 (10 μM) (Δ +0.2 ± 0.3%; N=6 vessels) (T_5_=1.223; p >0.05) (**Figs. 1Di-Dii**) and mimicked by application of a selective P2Y_2_ receptor agonist (PSB 1114; 200 nM) (Δ −2.3 ± 0.7%; N=11 vessels) (T_10_=3.460; p<0.010) (**Figs. 1Ei-Eii**). As a negative control, we confirmed that PSB 1114 (200 nM) minimally affects vascular tone in the cNTS (Δ −0.05 ± 0.09%; N=6 vessels; T_5_=0.3898; p>0.05) and ROb (Δ −0.07 ± 0.19%; N=7 vessels; T_6_=0.6076_;_ p>0.05) (**Figs. 1Ei-Eii**), where P2Y_2_ receptors were detected at low levels in both endothelial and smooth muscle cells (**Figs. 1A-B**). We also confirmed that the response of RTN arterioles to CO_2_/H^+^ was retained when neural action potentials were blocked by bath application of TTX (0.5 μM) (**Supplemental Figs. 1A-B**), indicating that CO_2_/H^+^ constriction of RTN arterioles differs from canonical neurovascular coupling both in terms of polarity and dependence (or lack thereof) on neuronal activity (21). Also, despite evidence that extracellular ATP can be rapidly metabolized to adenosine (a potent vasodilator including in the RTN (18)) the response of RTN arterioles to CO_2_/H^+^ was unaffected by 8-phenyltheophylline (8 PT; 10 μM) to block adenosine A1 receptors or sodium metatungstate (POM 1; 100 µM) to inhibit ectonucleotidase activity (**Supplemental Figs. 1A-B**). Furthermore, exposure to CO_2_/H^+^ has been shown to trigger prostaglandin E2 (PGE_2_) release that may contribute to RTN chemoreception by a mechanism involving EP3 receptors (10); therefore, we also tested for a role of PGE_2_/EP3 signaling in CO_2_/H^+^-dependent regulation of RTN arteriole tone. We found bath application of PGE_2_ (1 µM) decreased RTN arteriole tone by −2.7 ± 0.4% (T_9_ = 2.787, p = 0.0212). However, CO_2_/H^+^ constriction of RTN vessels was retained in the presence of L-798,106 (0.5 µM) to block EP3 receptors (−9.6 ± 4.0%, T_7_ = 2.675, p = 0.0318) (Supplemental Fig. 1A-B), suggesting PGE_2_/EP3 signaling is dispensable for CO_2_/H^+^ regulation of arteriole tone in the RTN.

**Table 1.** Respiratory parameters in control (αSm22^cre^ only and P2Y2 ^f/f^ only), P2Y_2_ cKO (αSm22^cre^::P2Y_2_ ^f/f^) and P2Y_2_ rescue animals two weeks after bilateral RTN injections of AAV2-Myh11p-eGFP-2A-mP2ry2. No significant differences were observed (p> 0.05).

| | αSm22<sup>cre</sup> only<br>P2Y<sub>2</sub><sup>f/f</sup> only | P2Y<sub>2</sub> cKO |
| --- | --- | --- |
| Frequency<br>(breaths/min) | 163.4 ± 7.1 | 149.8 ± 7.2 |
| Tidal Volume<br>(μLx10<sup>-3</sup>) | 6.8 ± 0.4 | 6.7 ± 0.5 |
| Minute Ventilation<br>(mL/min/kg) | 1.1 ± 0.1 | 1.0 ± 0.1 |
| Systolic blood pressure<br>(mm Hg) | 124.2 ± 0.9 | 123.0 ± 1.2 |
| Diastolic blood pressure<br>(mm Hg) | 85.5 ± 0.8 | 86.7 ± 1.4 |
| Heart rate<br>(beats/min) | 473.8 ± 3.0 | 465.0 ± 5.0 |
| VO<sub>2</sub> (mL/kg/hr) | 4886.5 ± 127.1 | 5067.0 ± 230.4 |
| VCO<sub>2</sub> (mL/kg/hr) | 4392.8 ± 101.9 | 4653.8 ± 204.5 |
| respiratory exchange<br>ratio (RER) | 0.89 ± 0.01 | 0.91 ± 0.01 |
| pH | 7.410 ± 0.005 | 7.406 ± 0.008 |
| pCO<sub>2</sub> (mmHg) | 40.1 ± 0.4 | 40.7 ± 0.6 |
| pO<sub>2</sub> (mmHg) | 88.3 ± 0.9 | 88.0 ± 0.8 |

Astrocytes are thought to contribute to CO_2_/H^+^ -induced vasodilation in the cortex (20) and vasoconstriction in the RTN (18). Consistent with this, we found that exposure to t-ACPD (50 µM), an mGluR agonist widely used to elicit Ca^2+^ elevations in astrocytes (20), caused constriction of RTN arterioles (Δ -5.9 ± 1.7%; N=8 vessels) and dilation of arterioles in the cNTS (Δ +7.6 ± 2.5%; N=8 vessels), ROb (Δ +6.8 ± 1.9%; N=8 vessels) and somatosensory cortex (Δ +2.9 ± 0.8%; N=5 vessels) (F_3,25_=10.18; p <0.0001) (**Fig. 1Fi-Fii**). These results show that CO_2_/H^+^-dependent regulation of vascular tone and roles of astrocytes in this process are fundamentally different in the RTN compared to other brain regions, and point to differential purinergic signaling mechanisms downstream of astrocyte activation.

To determine whether P2Y_2_ receptors in the RTN regulate CO_2_/H^+^ vascular reactivity *in vivo*, we measured the diameter of pial vessels on the ventral medullary surface (VMS) in the region of the RTN during exposure to high CO_2_ under control conditions (saline) and when P2Y_2_ receptor are blocked with AR-C118925. Consistent with our slice data, we found under saline control conditions that increasing end-expiratory CO_2_ to 9-10%, which corresponds with an arterial pH of 7.2 pH units (16), constricted VMS vessels by −7.8 ± 0.7 % (p=0.01, N = 7 animals) (**Fig. 2A**). However, when P2Y_2_-receptors are blocked by application of AR-C118925 (10 µM) to the VMS, increasing inspired CO_2_ resulted in a vasodilation of 5.2 ± 0.9% (**Fig. 2A**) (p=0.05; N = 7 animals). Thus, in the absence of functional P2Y_2_ receptors, RTN vessels respond to CO_2_/H^+^ in a manner similar to other brain regions. Also consistent with our slice data, we found that activation of P2Y_2_ receptors by application of PSB1114 (100 µM) to the VMS constricted vessels in the region of the RTN by −6.3 ± 0.9 % (p=0.05, N = 7 animals) (**Fig. 2B**).

**Figure 2.**
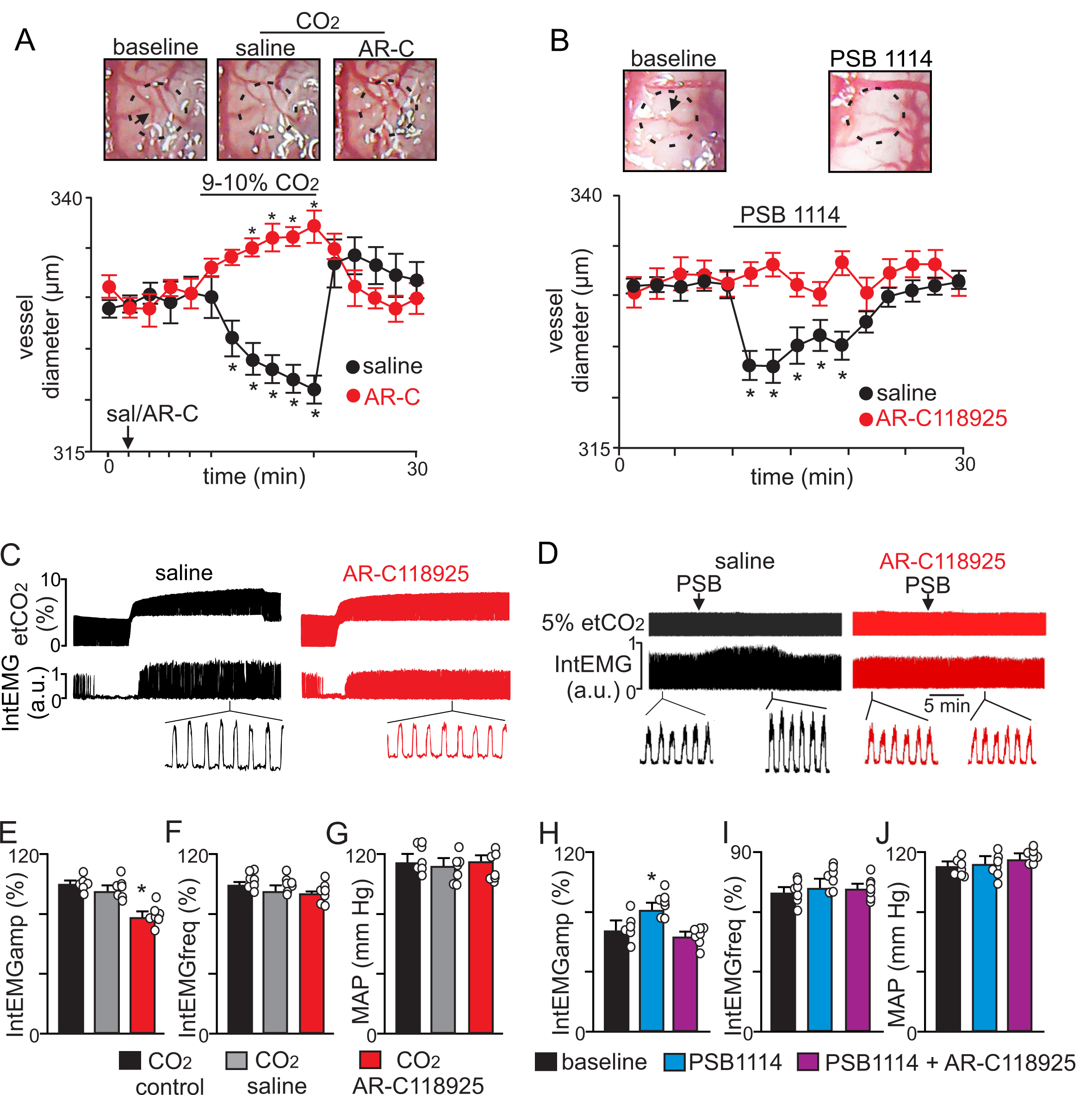
CO_2_ constricts pial vessels in the RTN region by a P2Y_2_ receptor-dependent mechanism to increase respiratory behavior in anesthetized mice. **A**, Images of RTN pial vessels and corresponding traces of vessel diameter (N=6 mice/condition) show that exposure to 9-10% CO_2_ decreased vessel diameter under control conditions (saline) but not when P2Y_2_ receptors were blocked with AR-C118925 (1 mM). **B**, Images of RTN pial vessels and corresponding traces vessel diameter show that application of a P2Y_2_ receptor agonist (PSB1114; 100 µM) caused a reversible constriction. **C-D**, Traces of external intercostal EMG (Int_EMG_) and end expiratory CO_2_ (etCO_2_) show that blocking CO_2_/H^+^-induced vasoconstriction by ventral surface application of AR-C118925 minimally affected respiratory activity at low etCO_2_ levels but blunted the ventilatory response to 9-10% CO_2_ (**C**). Conversely, at a constant etCO_2_ of 5% the application of PSB1114 to mimic CO_2_/H^+^ constriction increased respiratory output (**D**). **E-J**, summary data show (N=6 mice/condition) effects RTN application of saline, AR-C118925 or PSB1114 on intercostal EMG amplitude (**E, H**), frequency (**F**,**I**) and mean arterial pressure (MAP; **G, J**). *, different (RM-ANOVA followed by Bonferroni multiple-comparison test; *, p < 0.05). scale bar = 200 μm.

To correlate P2Y_2_-dependent vasoconstriction in the RTN region with respiratory behavior, we simultaneously measured systemic blood pressure and external intercostal electromyogram (Int_EMG_) activity (as a measure of respiratory activity) in urethane-anesthetized mice during exposure to low and high CO_2_ following VMS application of saline or AR-C118925. We found that VMS application of AR-C118925 (1 mM) minimally affected the CO_2_/H^+^ apneic threshold (3.3 ± 0.4% vs. saline: 3.1 ± 0.5% (p > 0.05; two-way RM ANOVA; N = 7). However at high levels of CO_2_ (9-10% etCO_2_) AR-C118925 decreased intercostal EMG amplitude by 23 ± 5% (F_2,74_ = 69.83; p=0.021; N = 7 animals) (**Figs. 2C, E**), but with no change in frequency (F_2,74_ = 1.09; p>0.05; N = 7 animals) (**Figs. 2C, F**). Conversely, VMS application of PSB1114 (100 µM) while holding etCO_2_ constant at 5% increased intercostal EMG amplitude by 20 ± 3% (F_2,74_ = 96.14; p=0.02; N = 7 animals) (**Figs. 2D, H**), again with no change in frequency (F_2,74_ = 0.79; p>0.05; N = 7 animals) (**Figs. 2D, I**). These treatments had negligible effects on systemic mean arterial pressure (MAP) (AR-C118925: 113 ± 5; PSB1114: 112 ± 5; saline: 114 ± 4 mmHg; F_2,74_ = 1.29; p>0.05; N = 7 animals) (**Figs. 2G, J**). These results indicate that P2Y_2_ dependent vasoconstriction in the RTN contributes to the drive to breathe.

If the mechanism underlying vascular control of RTN chemoreception involves maintenance of tissue CO_2_/H^+^ levels, then by this same logic, vasodilation in the cNTS and ROb should buffer tissue CO_2_/H^+^ and limit contributions of these regions to respiratory output. Such a braking mechanism may be important for stabilizing breathing since over-activation of the CO_2_/H^+^ chemoreflex is thought to cause unstable periodic breathing (5). To test this, we disrupted CO_2_/H^+^ dilation by injecting the vasoconstrictor U46619 (a thromboxane A2 receptor agonist) into the cNTS and ROb while measuring cardiorespiratory activity in anesthetized mice breathing a level of CO_2_ required to maintain respiratory activity (2-3% CO_2_). We found that injections of U46619 (1 μM) into the cNTS alone or together with the ROb resulted in unstable breathing as evidenced by frequent bouts of hyperventilation followed by apnea (**Fig. 3A**) (1.2 ± 0.6, vs. saline: 0.4 ± 0.2 apneas/min; p < 0.001) and increased breath to breath variability (**Figs. 3C-E**). Consistent with increased chemoreceptor drive, we found that application of U46619 into these regions lowered the CO_2_ apneic threshold from 3.2 ± 0.3% to 2.1 ± 0.1% (p < 0.05; two-way RM ANOVA; N = 7) (**Fig. 3F**). Also, consistent with previous work (41), we found that injection of U46619 (1 μM) into the cNTS also increased systemic blood pressure (127 ± 11, vs. saline: 98 ± 4 mmHg, (F_3,65_ = 77.62; p<0.01, data not shown), whereas, application of this drug into the ROb alone had negligible effects on cardiorespiratory output. Together, these results show that CO_2_/H^+^ induced vasoconstriction in the RTN serves to enhance chemoreceptor drive, while simultaneous CO_2_/H^+^ dependent dilation in the cNTS and possibly the ROb limits chemoreceptor activity to stabilize CO_2_/H^+^ stimulated respiratory drive.

**Figure 3.**
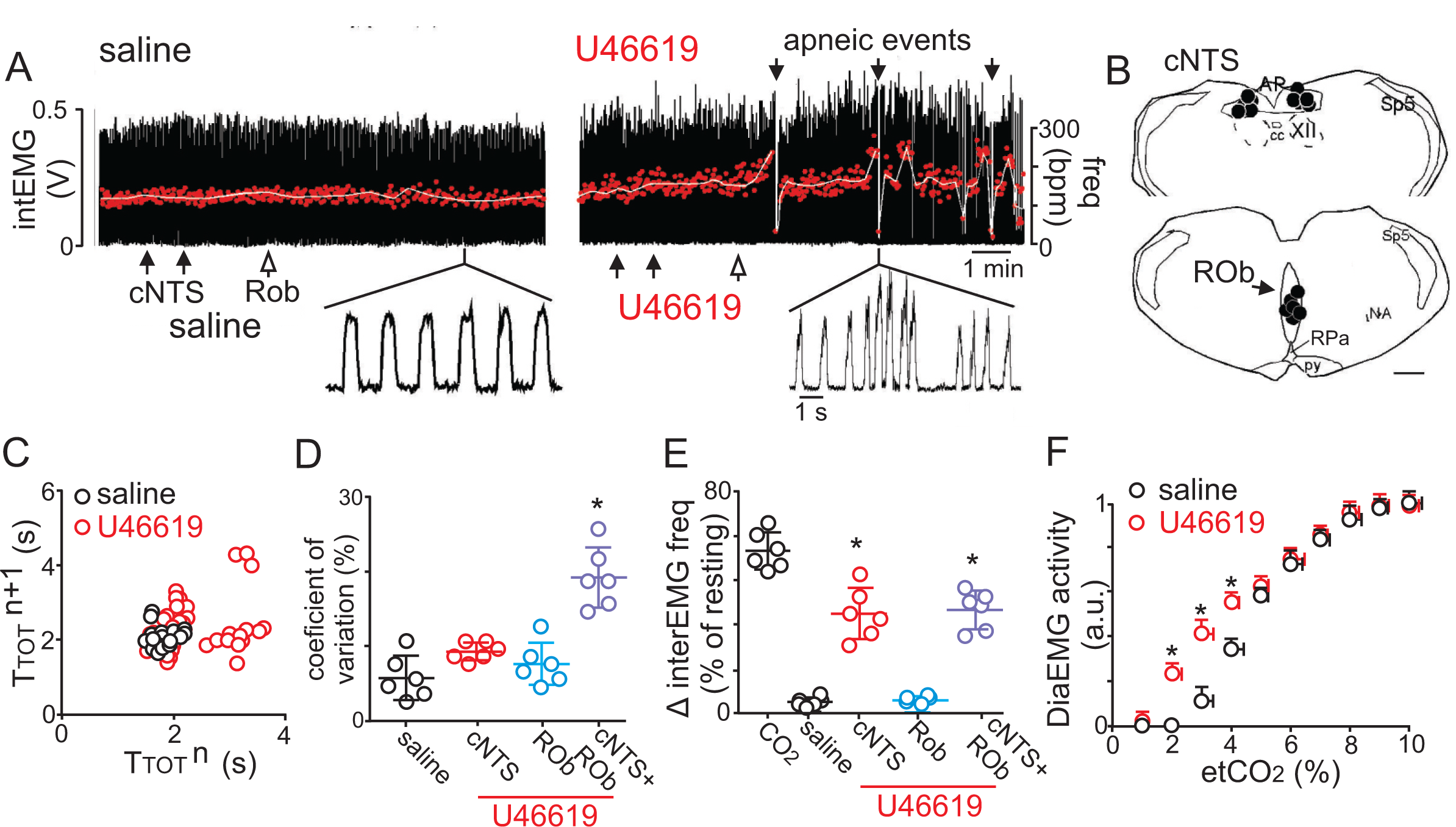
Disruption of CO_2_/H^+^ dilation in the cNTS and ROb causes unstable breathing and apnea. **A**, trace of external intercostal muscle EMG (Int_EMG_) activity shows respiratory activity of an anesthetized wild type mouse breathing 2.5% CO_2_ following injections of saline or U46619 (1 μM; 30 nL/region) into the cNTS and ROb. **B**, location of injections in the cNTS and ROb. **C**, Representative Poincaré plot (50 breaths) shows breath-to-breath (T_TOT_) interval variability following injections saline (black) or U46619 (red) conditions. **D-E**, Summary data (N=6 animals/group) shows effects of U46619 injections into the cNTS and ROb alone and in combination on the coefficient of variation of Int_EMG_ frequency (**C**) and Int_EMG_ frequency (**E**). **F**, Summary data show that injections of U46619 injections into the cNTS and ROb lowered the CO_2_ apneic threshold from 3.2 ± 0.3% to 2.1 ± 0.1% (N = 7 mice). *, difference in Int_EMG_ activity under control conditions (saline) vs. during U46119 into the NTS and/or ROb (RM-ANOVA followed by Bonferroni multiple-comparison test, p < 0.05). scale bar = 200 μm.

### Smooth muscle specific P2Y_2_ cKO mice have a blunted chemoreflex that can be rescued by targeted re-expression of P2Y_2_ in RTN smooth muscle cells

To definitively test contributions of P2Y_2_ receptors in vascular smooth muscle cells to respiratory activity, we created a smooth muscle cell specific P2Y_2_ knockout mouse (P2Y_2_ cKO maintained on a C57BL6/J background) by crossing αSm22^cre^ (JAX #: 017491) with P2Y_2_^f/f^ mice provided by Dr. Cheike Seye (Indiana Univ.). We confirmed that cre recombinase expression was restricted to vascular smooth muscle cells (**Fig. 4A**) and that P2Y_2_ receptor transcript was not detectable in RTN smooth muscle cells from P2Y_2_ cKO mice (**Fig. 4A**). Each genotype (αSm22^cre^ only, P2Y_2_^f/f^ only and P2Y_2_ cKO mice) was obtained at the expected ratios and gross motor activity, metabolic activity, blood gases, heart rate and blood pressure were all similar between genotypes and sexes (**Table 1, Supplemental Fig. 2**). Therefore, results from male and female mice of each genotype were pooled. To determine whether P2Y_2_ cKO mice exhibit respiratory problems, we used whole-body plethysmography to measure baseline breathing and the ventilatory response to CO_2_ in conscious unrestrained adult mice (mixed sex). Consistent with the possibly that smooth muscle P2Y_2_ receptor-mediated vasoconstriction in the RTN augments the ventilatory response to CO_2_, we found that P2Y_2_ cKO mice exhibit normal respiratory activity under room air conditions **(Table 1)** but pronounced hypoventilation during exposure to graded increases in CO_2_ (balance O_2_ to minimize input from peripheral chemoreceptors). For example, minute ventilation – the product of frequency and tidal volume – of P2Y_2_ cKO mice at 5% and 7% CO_2_ was 28% and 35% smaller than control counterparts (**Figs. 4C-F, Table 2**). This chemoreceptor deficit involves a diminished capacity to increase both respiratory frequency (**Fig. 4D**) and tidal volume (**Fig. 4E**) during exposure to CO_2_ (**Table 2**). Similar results were obtained in P2Y_2_ cKO mice generated using a different smooth muscle cre line (smMHC^cre/eGFP^) (**Supplemental Fig. 3**). Also, smooth muscle P2Y_2_ cKO mice showed a normal ventilatory response to hypoxia (10% O_2_; balance N_2_, T_7_=0.1089, p>0.05) (**Supplemental Fig. 4**). We confirmed *in vitro* that RTN arterioles in slices from P2Y_2_ cKO do not respond to PBS 1114 (0.4 ± 0.1%, T_7_ = 1661, p > 0.05, N=8 vessels). Interestingly, we also found that RTN arterioles in slices from P2Y_2_ cKO mice fail to constrict in response to CO_2_/H^+^ or tACPD, but rather respond in a manner similar to arterioles in other brain regions. For example, both CO_2_/H^+^ and tACPD dilated RTN (CO_2_/H^+^: 4.7 ±0.2%, T_9_ = 3.312, p = 0.009, N=10 vessels; tACPD: 3.2 ± 0.7%, T_7_ = 2.864, p = 0.035, N = 8 vessels) and cortical (CO_2_/H^+^: 7.3 ± 1.5%, T_6_=4.307, p=0.0051, N=7 vessels; tACPD: 6.7 ± 0.9%, T_6_=2.739, p=0.0338, N=7 vessels) arterioles in slices from P2Y_2_ cKO mice (**Supplemental Fig. 5A-B**). This suggests CO_2_/H^+^ induced vasodilation is a general phenomenon mediated by undetermined mechanisms, but in the RTN is response is countered by smooth muscle P2Y_2_ receptor-mediated constriction.

**Table 2.**
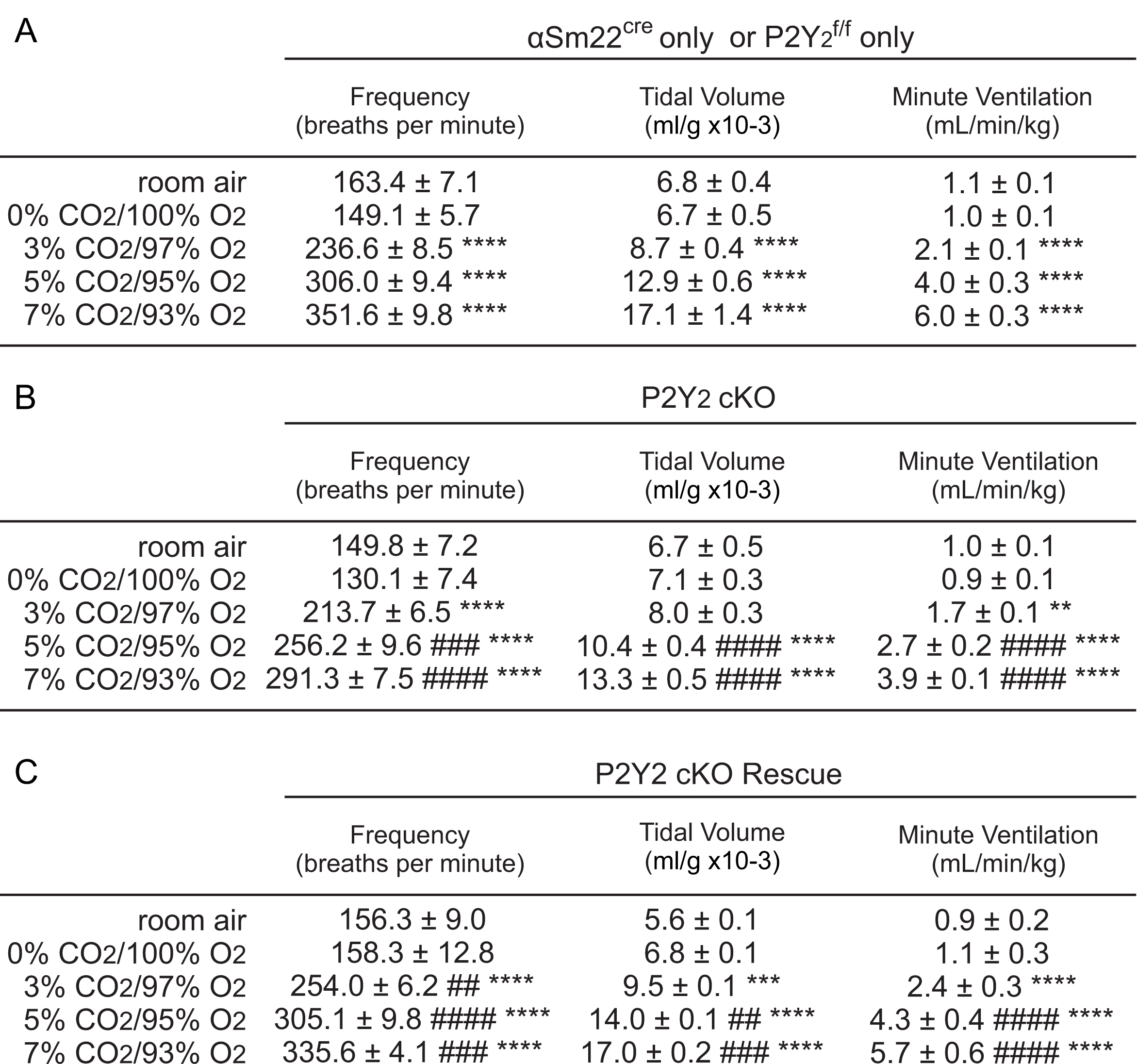
Cardiorespiratory and metabolic parameters under room air conditions in control and P2Y_2_ cKO mice. *, difference from 0% CO_2_; ^#^, differences between genotypes under an experimental condition (two-way ANOVA with Tukey’s multiple comparison test). Two symbols = p < 0.01, three symbols = p < 0.001, four symbols = p < 0.0001.

**Figure 4.**
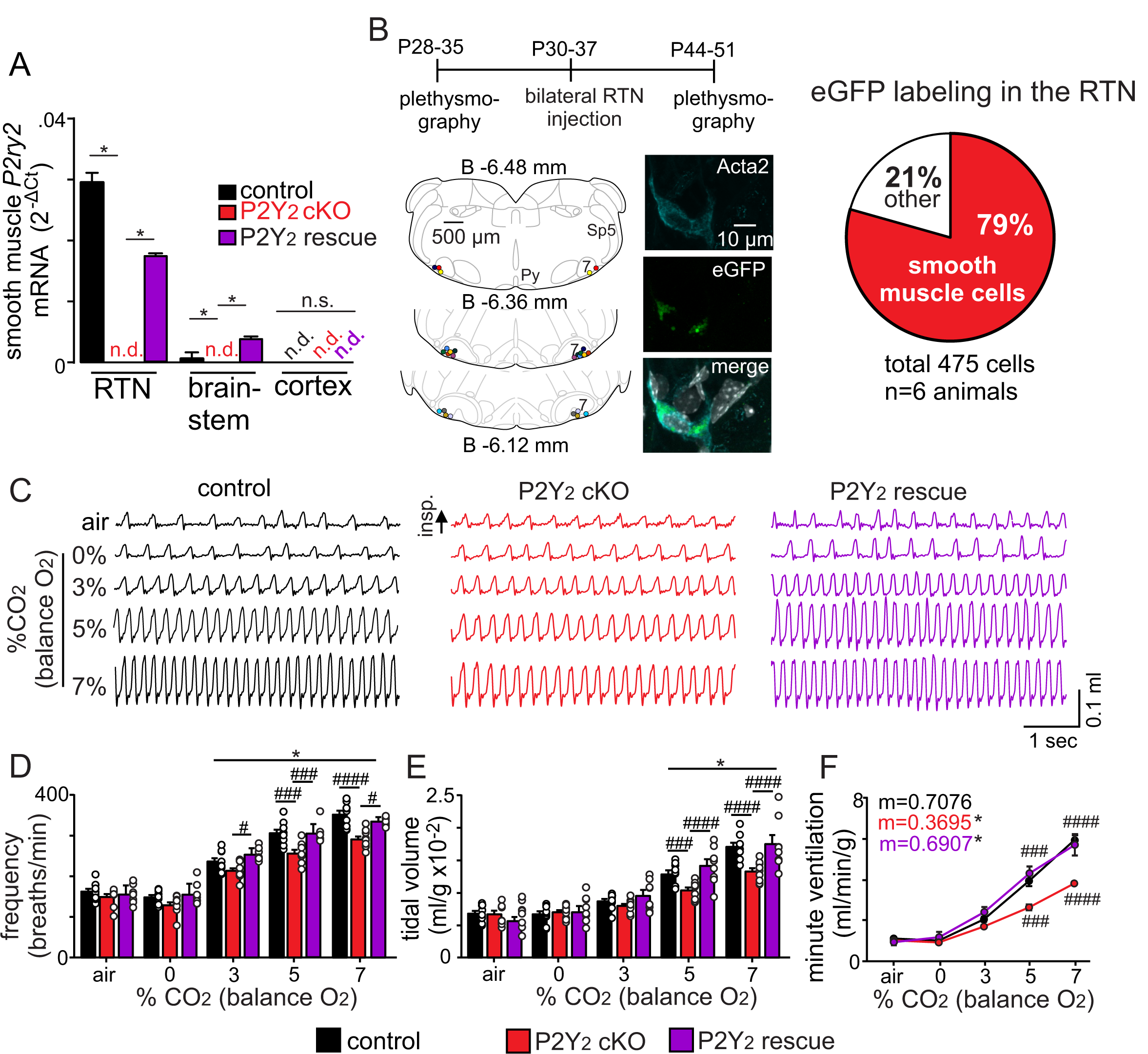
Smooth muscle P2Y_2_ cKO mice show a blunted CO_2_ chemoreflex that can be rescued by re-expression of P2Y_2_ only in RTN smooth muscle cells. **A**, P2Y_2_ transcript was detected in RTN, brainstem and cortical smooth muscle cells isolated from control mice (αSm22^Cre^::TdTomato), P2Y_2_ cKO mice (αSm22^Cre^::P2Y_2_^f/f^::TdTomato), and P2Y_2_ rescue mice (P2Y_2_ cKO animals that received bilateral RTN injections of AAV2-Myh11p-eGFP-2A-mP2ry2). P2Y_2_ transcript was not detected (n.d.) in smooth muscle cells from P2Y_2_ cKO mice (N=3 runs/9 animals). P2Y_2_ was also not detected in cortical smooth muscle cells from either genotype. Conversely, RTN (p=0.0073) and brainstem (p=0.0073) but not cortical (p>0.05) smooth muscle cells from P2Y_2_ rescue mice show increased P2Y_2_ transcript levels compared to P2Y_2_ cKO but not to the same level as cells from control mice (p=0.0219) (ANOVA on ranks followed by Dunn multiple comparison test). **B** Left, computer-assisted plots show the center of all bilateral AAV2-Myh11p-eGFP-2A-mP2ry2 injections; each matching color pair of dots corresponds to one animal (N=13 animals). Approximate millimeters behind bregma (33) is indicated by numbers next to each section. Right, two weeks after injections we confirmed that ∼80% of RTN smooth muscle α2 actin (Acta2)-immunoreactive cells were also GFP^+^(inset). **C-F**, representative traces of respiratory activity (**C**) and summary data show that smooth muscle-specific P2Y_2_ KO mice (αSm22^Cre^::P2Y_2_^f/f^) breathe normally under room air conditions but fail to increase respiratory frequency (**C**) or tidal volume (**D**) during exposure to CO_2_, thus resulting in diminished minute ventilation at 5-7% CO_2_ (**E**). Re-expression of P2Y_2_ in only RTN smooth muscle cells rescued of the ventilatory response to CO_2_. Note that αSm22^cre^ only and P2Y_2_^f/f^ only control mice showed similar baseline breathing and responses to CO_2_ and so were pooled. *, different from 0% CO_2_ in condition as assessed by Tukey’s post-hoc multiple comparison test. ^####^, different between genotypes (two-way ANOVA with Tukey’s multiple comparison test, p<0.0001).

We next tested whether targeted re-expression of P2Y_2_ in RTN smooth muscle cells would rescue the blunted chemoreflex in P2Y_2_ cKO mice. An adeno-associated virus (AAV2-Myh11p-eGFP-2A-mP2ry2, Vector Biolabs) was injected bilaterally into the RTN of adult P2Y_2_ cKO (αSm22^cre^/P2Y_2_^f/f^) to drive P2Y_2_ expression in smooth muscle cells. Two weeks after virus injection into the RTN region, we confirmed that ∼80% of smooth muscle α2 actin (Acta2)-immunoreactive cells in the RTN are also GFP^+^ (**Fig. 4B, Supplemental Fig. 6**). Some background eGFP labeling was observed, but not associated with DAPI labeling (**Supplemental Fig. 6aii)**. We also confirmed RTN smooth muscle cells from P2Y_2_ rescue mice show increased P2Y_2_ transcript levels compared to P2Y_2_ cKO (p=0.0073) but not to the same level as cells from control mice (p = 0.0219) (**Fig. 4A**). Importantly, re-expression of P2Y_2_ receptors in RTN smooth muscle cells resulted in a full rescue of the minute ventilatory response to CO_2_ (**Fig. 4C-F, Table 2**) (F_1,7_=17.75, p=0.004, N= 13 animals). Conversely, bilateral RTN injections of control virus (AAV2-Myh11p-eGFP, Vector Biolabs) minimally affected respiratory parameters of interest in P2Y_2_ cKO mice (**Supplemental Fig. 3**). Furthermore, re-expression of P2Y_2_ receptors in RTN smooth muscle cells also rescues CO_2_/H^+^ vascular reactivity. Specifically, we found in slices from P2Y_2_ cKO mice, that RTN arterioles transfected with AAV showed a robust constriction in response to CO_2_ (−19.9 ± 6.9%, p=0.0421), tACPD (−6.2 ± 1.9%, p=0.0092) and PSB1114 (−9.3 ± 4.9%, p=0.0154) (**Supplemental Fig. 5**). These results identify the first vascular element of respiratory control by showing that P2Y_2_ receptors in RTN vascular smooth muscle cells are required for the normal ventilatory response to CO_2_

## Discussion

These results identify P2Y_2_ receptors in RTN smooth muscle cells as a novel vascular element of respiratory chemoreception. We show that CO_2_/H^+^ constriction by activation of smooth muscle P2Y_2_ receptors is unique to the RTN and is required for the normal ventilatory response to CO_2_, whereas simultaneous CO_2_/H^+^ vasodilation in other chemoreceptor regions like the cNTS and ROb may serve to stabilizes breathing. Considering these regions sense changes in CO_2_/H^+^ to regulate breathing, and since vasoconstriction and dilation are expected to increase and decrease tissue CO_2_/H^+^ levels respectively, we speculate that differential CO_2_/H^+^ vascular reactivity across multiple chemoreceptor regions serves to fine tune respiratory drive and stability during exposure to high CO_2_. Understanding this novel mechanism may lead to new treatment options for breathing problems, particularly those associated with cardiovascular disease.

### Experimental Limitations

There are several limitations of this work that should be recognized. First, our *in vitro* experiments utilized the slice preparation because it allows for easy visualization of vessels in each region of interest and pharmacological manipulation of candidate mechanisms independent of potential confounding effects of blood pressure on myogenic tone. However, because blood vessels in brain slice have limited myogenic tone these experiments were performed in the presence of U-46619 to pre-constrict vessels ∼30% (2). We also used a large stimulus (15% CO_2_) to characterize CO_2_/H^+^ vascular reactivity *in vitro*. For these reasons, we also confirmed *in vivo* in the absence of U-46691 that more physiological levels of CO_2_ (9-10%) constricted arterioles in the RTN by a P2Y_2_-dependent mechanism. Second, we and others (41) showed that administration of U46619 into the NTS increased blood pressure. Although baroreceptor activation has a fairly mild effect on respiratory activity (28), it remains possible that the baroreflex contributed to respiratory instability observed during this experimental manipulation. Furthermore, neurons and astrocytes may express thromboxane A2 receptors (12, 30) so it is possible respiratory instability caused by injections of U46619 into the cNTS and ROb may involve direct neural or astrocyte activation. Third, CO_2_/H^+^ typically increases minute ventilation by increasing both rate and depth of breathing (36, 37), yet ventral surface application of drugs to manipulate P2Y_2_ receptors only affected tidal volume (**Figs. 2E-J**). Therefore, it is possible other nearby respiratory centers were affected in this experiment. However, this issue is mitigated by more targeted genetic approaches showing that deletion of P2Y_2_ from smooth muscle cells blunted the rate and depth of respiratory responses to CO_2_, and re-expression of P2Y_2_ receptors only in RTN smooth muscle cells fully rescued the CO_2_/H^+^ chemoreflex.

### Mechanisms contributing to vascular control of breathing

Blood flow in the brain is controlled by several mechanisms including neural activity, which leads to vasodilation and increased blood flow by a process termed neurovascular coupling (21). Autoregulation is a mechanism that maintains cerebral blood flow in response to changes in perfusion pressure (32). For example, myogenic tone increases with increased perfusion pressure and relaxes with decreased pressure. CO_2_/H^+^ also functions as a potent vasodilator in most brain regions (19), and is perhaps the most potent regulator of cerebrovascular tone since high CO_2_/H^+^ can impair both autoregulation (31, 34) and neurovascular coupling (25, 26). Since CO_2_/H^+^ is a waste product of metabolism, CO_2_/H^+^ vascular reactivity provides a means of matching local blood flow with tissue metabolic needs. However, at the level of the RTN this need appears secondary to chemoreceptor regulation of respiratory activity. For example, we (18) and others (40) showed that exposure to high CO_2_/H^+^ constricted RTN arterioles, and disruption of this response blunted the ventilatory response to CO_2_.

The cellular and transmitter basis for CO_2_/H^+^ induced constriction of RTN arterioles appears to involve CO_2_/H^+^-evoked ATP release from local astrocytes. For example, previous work showed that i) CO_2_/H^+^ caused discrete release of ATP near the RTN region (15); ii) astrocytes in the RTN show Ca^2+^ responses to mild acidification (14); and iii) activation of astrocytes with tACPD, an mGluR agonist widely used to elicit Ca^2+^ elevations in astrocytes (20), constricted RTN arterioles by a purinergic-dependent mechanism (18) (**Fig. 2B**), whereas application of an ATP receptor blocker to the ventral surface blunted CO_2_/H^+^ induced vasoconstriction and the ventilatory response to CO_2_ (18). Here, we extend this work by showing that CO_2_/H^+^ vascular responses in the RTN are opposite to other chemoreceptor regions, and we identify P2Y_2_ receptors in vascular smooth muscle cells as requisite determinants of RTN CO_2_/H^+^ vascular reactivity and chemoreception. For example, P2Y_2_ is a metabotropic receptor that couples with Gq, Go and G12 proteins (8); activation of this receptor in smooth muscle is associated with vasoconstriction (3, 23), whereas its activation in endothelial cells favors nitric oxide-mediated vasodilation (27). Consistent with a role in vasoconstriction, we found that P2Y_2_ transcript was expressed at higher than control levels in RTN smooth muscle cells and at lower than control levels in RTN endothelial cells (**Figs. 1A-B**). Furthermore, pharmacological experiments show *in vitro* (**Figs. 1C-E**) and in anesthetized mice (**Figs. 2A-B**) that CO_2_/H^+^-induced constriction of RTN arterioles was eliminated by blocking P2Y_2_ receptors with AR-C118925 and mimicked by a selective P2Y_2_ receptor agonist (PSB 1114). We also show in awake mice that genetic deletion of P2Y_2_ from smooth muscle cells blunted the ventilatory response to CO_2_, and re-expression of P2Y_2_ receptors only in RTN smooth muscle cells fully rescued the CO_2_/H^+^ chemoreflex (**Figs. 4A-F**). These results establish P2Y_2_ receptors in RTN smooth muscle cells as requisite determinants of respiratory chemoreception.

It should be noted that other cell types and signaling pathways may also contribute to RTN vascular reactivity. For example, endothelial cells including those in the RTN, express H^+^ activated GPR4 receptors that when activated facilitates release of vasoactive effectors including thromboxane to mediate vasoconstriction (40). Consistent with this, disruption of GPR4 signaling blunted CO_2_/H^+^ vascular reactivity in the RTN and the ventilatory response to CO_2_ (40). Although untested, it is also possible RTN endothelial cells release other vasoactive signals including ATP, and so may contribute to CO_2_/H^+^-evoked purinergic-dependent constriction of arterioles in this region. Prostaglandin E2 (PGE_2_) is also thought to contribute to CO_2_/H^+^ vascular reactivity. For example, in the cortex CO_2_/H^+^ facilitates Ca^2+^ oscillations in astrocytes (14, 20) leading to enhanced arachidonic acid metabolism and release of PGE_2_, which triggered arteriole dilation by activation of prostaglandin (EP1) receptors (20). In the RTN, CO_2_/H^+^ also causes PGE_2_ release most likely from astrocytes (11) and contributes to RTN chemoreception by a mechanism involving EP3 receptors (10). However, selective blockade of EP3 receptors minimally affected CO_2_/H^+^ constriction of RTN arterioles (**Supplemental Fig. 1B**), suggesting PGE_2_/EP3 signaling contributes to RTN chemoreception directly by modulation of chemosensitive neurons.

In contrast to the RTN, we also found that CO_2_/H^+^ (**Fig. 1C**) and astrocyte activation by bath application of t-ACPD (**Fig. 1F**) dilatated arterioles in other chemoreceptor regions including the cNTS and ROb. This suggests that CO_2_/H^+^-dependent regulation of vascular tone and roles of astrocytes in this process are fundamentally different in the RTN compared to other chemoreceptor regions. Considering vasoconstriction and dilation are expected to increase and decrease tissue CO_2_/H^+^ levels respectively, we propose that CO_2_/H^+^ constriction in the RTN supports the drive to breathe by ensuring tissue acidification is maintained to stimulate chemoreceptors in this region, whereas CO_2_/H^+^ dilation in the cNTS and ROb provides a means of buffering tissue H^+^ and chemoreceptor activity to maintain respiratory stability. Conversely, if CO_2_/H^+^ were to cause vasoconstriction across multiple chemoreceptor regions simultaneously, this may exaggerate the ventilatory response to CO_2_ to the point of causing unstable periodic breathing (5). Indeed, this is thought to be the cause of unstable breathing in patients with global deficits in CO_2_/H^+^ vascular reactivity due to congestive heart failure or cerebrovascular disease (4, 5). Consistent with this, we show that disruption of CO_2_/H^+^ dilatation in the cNTS and ROb by application of the vasoconstrictor U46619 while leaving RTN CO_2_/H^+^ constriction unperturbed, increased chemoreceptor drive as evidenced by a decrease in the CO_2_/H^+^ apneic threshold and caused unstable periodic breathing during exposure to higher levels of CO_2_/H^+^ (**Fig. 3**). However, further work is needed to understand whether and how differential CO_2_/H^+^ vascular reactivity in these chemoreceptor regions contributes to breathing problems in disease states.

## ACKNOWLEDGEMENTS

The authors thank Drs. Douglas Bayliss (Univ. Virginia) and Akiko Nishiyama (Univ. Connecticut) for their comments on the manuscript. This work was supported by funds from the National Institutes of Health Grants HL104101 (DKM), HL137094 (DKM), NS099887 (DKM) R01NS110656 (MTN), R35HL140027 (MTN) and F31HL142227 (CMC). Additional funds were also provided by the American Heart Association (17SDG33670237 and 19IPLOI34660108 to TAL), the São Paulo Research Foundation (FAPESP; grants: 2019/01236-4 to ACT; 2015/23376-1 to TSM) and the Conselho Nacional de Desenvolvimento Científico e Tecnológico (CNPq; grant: 408647/2018–3 to ACT). CNPq fellowships were awarded to ACT (302288/2019-8) and to TSM (302334/2019-0). This study was also financed by the Coordenação de Aperfeiçoamento de Pessoal de Nível Superior - Brasil (CAPES) - Financial Code 001 and by Serrapilheira Institute (Serra-1812-26431 to ACT). This work was also supported by Fondation Leducq (Transatlantic Network of Excellence on the Pathogenesis of Small Vessel Disease of the Brain) (MTN), the European Union (Horizon 2020 Research and Innovation Programme SVDs@target under the grant agreement n° 666881) (MTN) and the Henry M. Jackson Foundation for the Advancement of Military Medicine, HU0001-18-2-0016 (MTN).

## AUTHOR CONTRIBUTIONS

CMC: experimental design; collection and analysis of data; revising the manuscript; final approval of the manuscript.

TSM: experimental design; collection and analysis of data; revising the manuscript; final approval of the manuscript.

ACT: collection and analysis of data; revising the manuscript; final approval of the manuscript MTN: experimental design; final approval of the manuscript.

TAL: experimental design; collection and analysis of data; revising the manuscript; final approval of the manuscript.

DKM: experimental design; data analysis; drafting the manuscript; revising the manuscript, final approval of the manuscript.

## MATERIALS AND METHODS

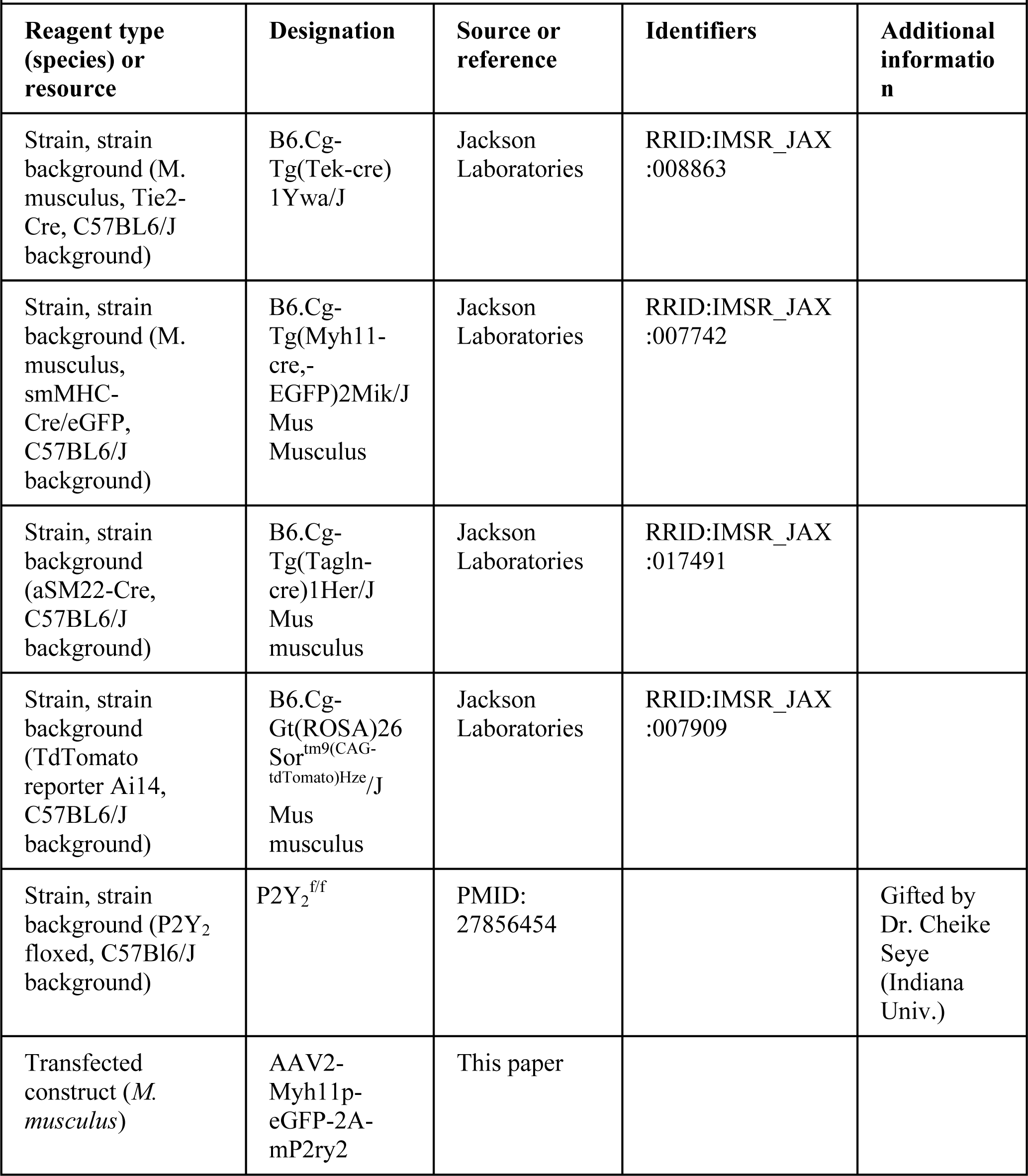
Key Resources Table.

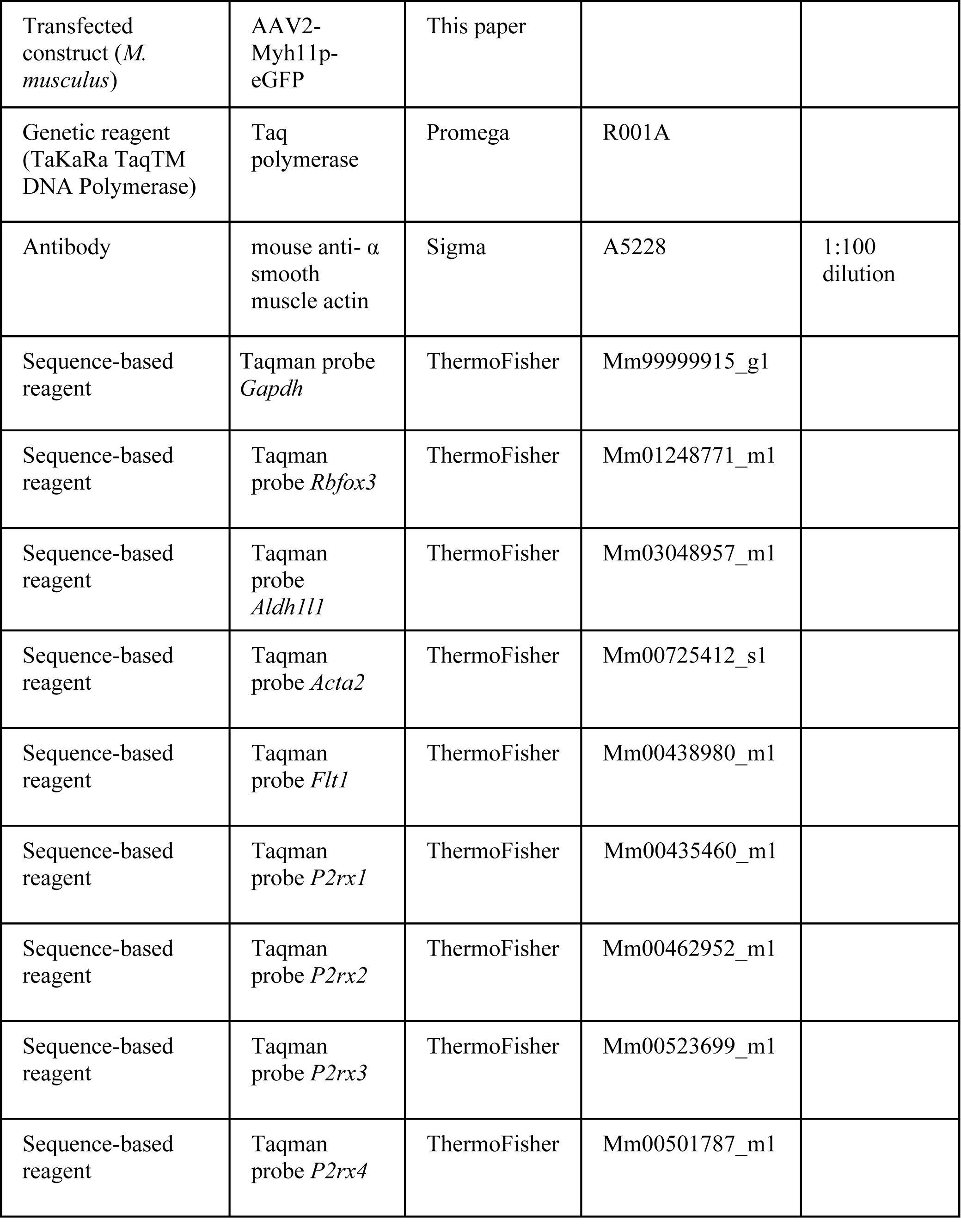

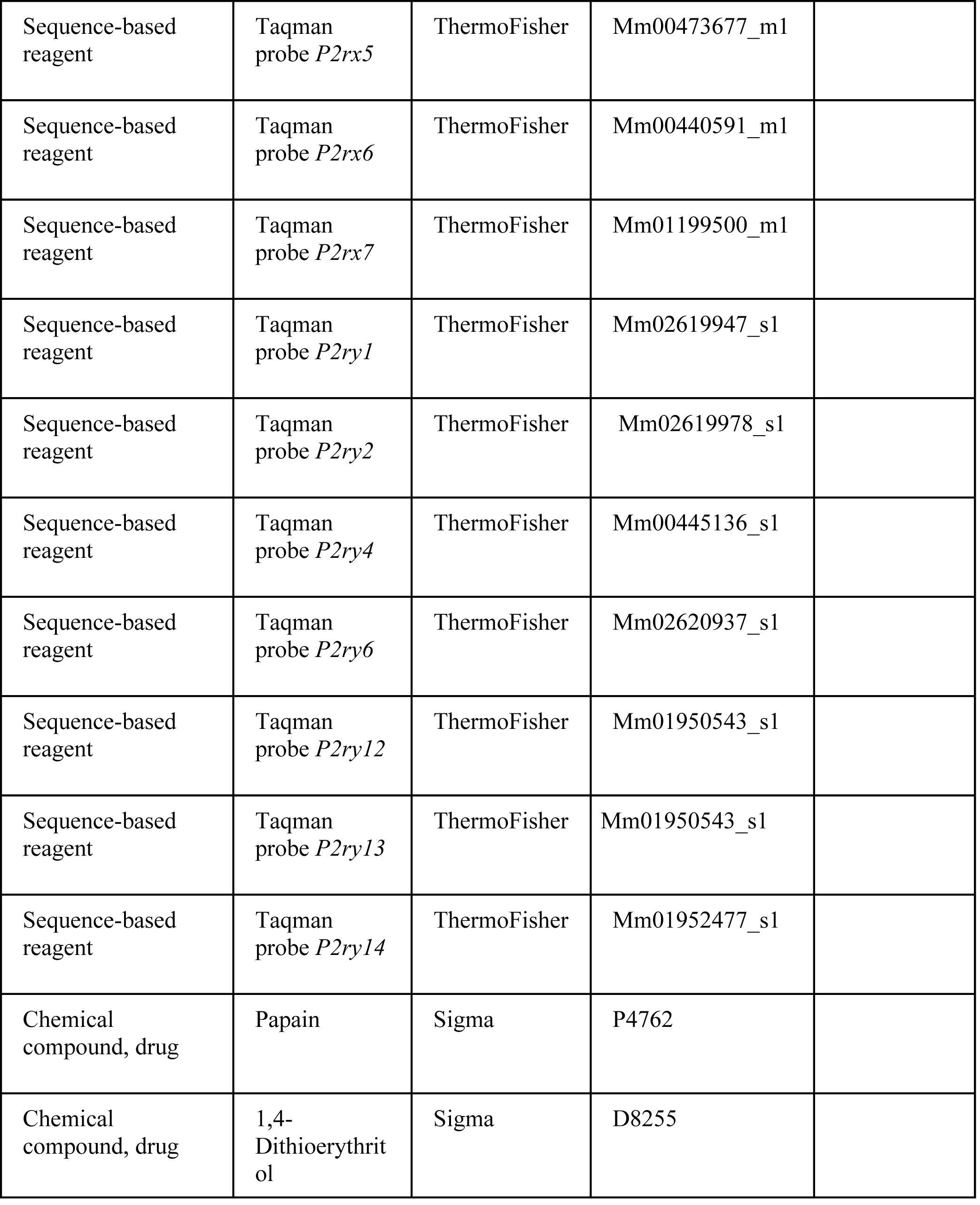

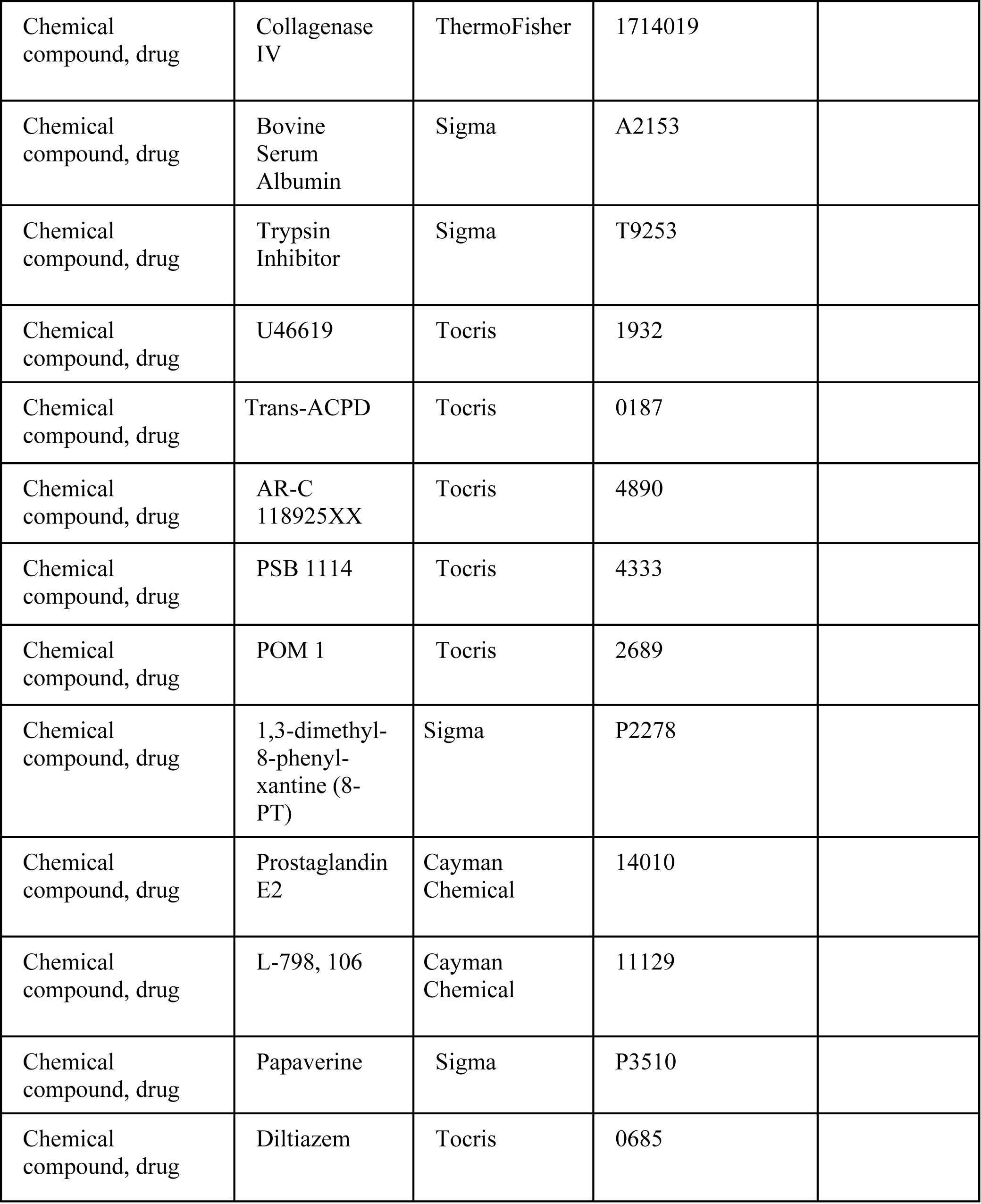

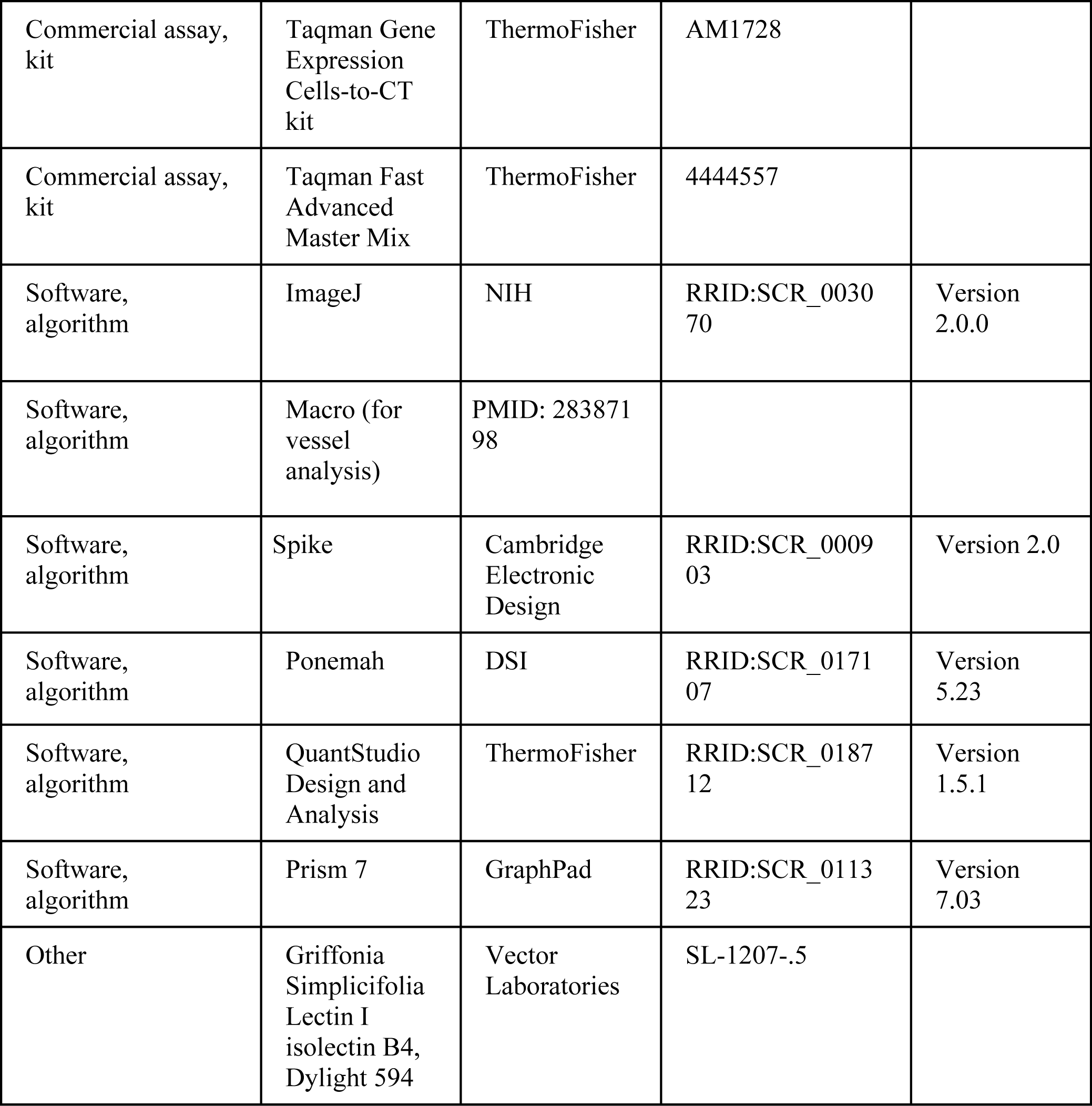

### Animals

All procedures were performed in accordance with National Institutes of Health and University of Connecticut Animal Care and Use Guidelines. All experiments used mixed sex C57BL6/J animals at least 3 weeks of age that housed in a 12:12 light dark cycle with normal chow *ad libitum*. The following transgenic cre mouse lines were used to perform single cell isolation, FACS, and pooled qPCR: smMHC^Cre/eGFP^ (Jax Stock #: 007742) and Tie2-Cre/TdTomato (Jax Stock #: 008863, 007909), and αSm22-Cre/TdTomato (Jax Stock #: 017491, 007909). αSm22-Cre and smMHC^Cre/eGFP^ mice were crossed with P2Y_2_ floxed mice (gifted by Dr. Cheike Seye at Indiana University) to generate double floxed smooth muscle specific P2Y_2_ cKO mice for *in vitro* and *in vivo* experimentation.

### *In vitro* arteriole slice recordings

#### Brainstem and Cortical Slice Preparation

Animals (P21 and older) were decapitated under isoflurane anesthesia and transverse brainstem slices (150 μm) were prepared using a vibratome in ice-cold substituted artificial cerebrospinal fluid (aCSF) solution containing (in mM): 130 NaCl, 3 KCl, 2 MgCl_2_, 2 CaCl_2_, 1.25 NaH_2_PO_4_, 26 NaHCO_3_, 10 glucose. 0.4 mM L-ascorbic acid was added into aCSF while slicing. Slices were incubated for 30 minutes at 37°C and subsequently at room temperature in aCSF, equilibrated with a 5% CO_2_-95% O_2_ gas mixture. Prior to imaging, each slice was incubated for thirty-five minutes with 6 mg/mL DyLight 594 Isolectin B4 conjugate (Vector Labs) to label vascular endothelium as previously described(18).

#### Imaging arterioles in vitro

Individual brain slices containing either the caudal NTS (cNTS), raphe obscurus (ROb), RTN, or somatosensory cortex were transferred to a Plexiglas recording chamber mounted on a fixed-stage upright fluorescent microscope (Zeiss Axioskop FS) and perfused with 37°C aCSF bubbled with a 5% CO_2_-21% O_2_ (balance N_2_). Hypercapnic solutions were made by allowing aCSF to equilibrate with a 15% CO_2_-21% O_2_ (balance N_2_). Arterioles were identified based on clear evidence of vasomotion under IR-DIC microscopy and bulky fluorescent labeling that indicates a thick layer of tightly wrapped smooth muscle surrounding the vessel lumen. Precapillary arterioles (as indicated by more sporadic, less uniform smooth muscle cell layer and thinner luminal diameter) were excluded from experimentation. All arterioles selected for experimentation had a luminal diameter between 8-50 µm. It should be noted that most, if not all, of the cNTS arterioles were small (average diameter was 9 µm) and ROb vasculature did not vary more than 10 µm from the midline of the slice. RTN vessels were located within 200 µm of the ventral surface and below the caudal end of the facial motor nucleus. Neocortical vessels were located in layers 1-5.

For an experiment, fluorescent images were acquired at a rate of 1 frame every 20 seconds using a 40X water objective lens, a Clara CCD Andor camera, and the NIS Advanced Research software suite (Nikon). To induce a partially constricted state in arterioles, we continuously bath applied 125 nM of a thromboxane A2 receptor agonist (U46619). U46619 has been previously shown to induce a 20-30% vasoconstriction at a concentration of 125 nM, enabling the vessel to either dilate or constrict further under experimental pharmacological studies (13). As previously described (18), we assessed arteriole viability at the end of each experiment by inducing a large vasoconstriction with a 60 mM K^+^ solution and then a large vasodilation with a Ca^+2^ free solution containing EGTA (5 mM), a phosphodiesterase inhibitor (papaverine, 200 mM), and an L-type Ca^+2^ channel blocker (diltiazem, 50 mM). Only one vessel was recorded per experiment and slice. Any vessel that did not respond to these solutions were excluded in data analysis. The list of all drugs and concentrations used for *in vitro* and *in vivo* experimentation are detailed in **Key Resource Table** and text where appropriate.

#### Image Analysis

Vessel diameter was determined using ImageJ. All images were calibrated, and pixel distance was converted to millimeters. Data files underwent StackReg (Biomedical Imaging Group) to stack each image over time and then three linear region of interests (ROI) were drawn orthogonal to the vessel. A macro (available at https://github.com/omsai/blood-vessel-diameter [Nanda, 2017]) was used to determine peak to peak distance using fluorescence intensity profile plots for all slices of the data file.

### Single cell isolation and qRT-PCR

At least three and no greater than eight, animals (postnatal days P21-P40) were used of either of the following genotypes: smMHC^Cre/eGFP^, Tie2-Cre/TdTomato, αSm22-Cre/TdTomato, or αSm22-Cre/TdTomato/P2Y_2_^fl/fl^. Animals were euthanized under isoflurane anesthesia and brainstem slices were prepared using a vibratome in ice cold, high sucrose slicing solution containing (in mM): 87 NaCl, 75 sucrose, 25 glucose, 25 NaHCO_3_, 1.25 NaH_2_PO_4_, 2.5 KCl, 7.5 MgCl_2_, 0.5 mM CaCl_2_ and 5 L-ascorbic acid. Slicing solution was equilibrated with a 5% CO_2_-95% O_2_ gas mixture before use. Transverse brainstem slices (400 μm thick) were prepared and then immediately enzymatically treated at 34°C with an initial incubation in a papain (6 mg/mL, Sigma) and dithioerythritol (10 mg/mL, ThermoFisher) mixture for 30 minutes in sucrose dissociation solution containing (in mM): 185 sucrose, 10 glucose, 30 Na_2_SO_4_, 2 K_2_SO_4_, 10 HEPES, 0.5 CaCl_2_, 6 MgCl_2_, 5 L-ascorbic acid, pH 7.4, 320 mOsm, followed by a collagenase IV (10 mg/mL, ThermoFisher) enzyme treatment for 6 minutes. After enzyme incubation, slices were washed three times in cold dissociation solution and then transferred to an enzyme inhibitor mix containing trypsin inhibitor (10 mg/mL, Sigma), bovine serum albumin (BSA, 10 mg/mL, Sigma), and sodium nitroprusside (SNP, 2 mg/mL, ThermoFisher) in cold sucrose dissociation solution. Shortly thereafter, slices were transferred to a glass Petri dish on ice. Using a plastic transfer pipette and a 15 blade scalpel, each region of interest was microdissected out of the slices and manually separated into sterile microcentrifuge tubes. The control samples were made up of two slices: one slice with the NTS removed and the other with the ROb and RTN removed. Once microdissection is completed, the tissue chunks were warmed to 34°C for 10 minutes before trituration. A single cell suspension was achieved by trituration using a 25G and 30G needle sequentially, attached to a 3 mL syringe. Samples were triturated for an average of 5 minutes. Immediately after, the samples were placed back on ice and filtered through a 30-micron filter (Miltenyi Biotech) into round bottom polystyrene tube for fluorescence-activated cell sorting (FACS).

#### Florescence-activated cell sorting (FACS)

All cell types of interest were sorted on a BD FACSAria II Cell Sorter (UConn COR^2^E Facility, Storrs, CT) equipped with 407 nm, 488 nm, and 607 nm excitation lasers. Five minutes before sorting, 5 µL of 100 ng/mL DAPI was added to each sample. Cells were gated based on scatter (forward and side), for singlets, and for absence of DAPI. Finally, cells were gated either to TdTomato or GFP fluorescence and sorted by 4-way purity into a sterile 96-well plate containing 5 µL of sterile PBS per sample. Between 100 and 500 cells were sorted per sample in any experiment and were processed immediately following FACS. To control for RNA contamination in FACS droplets, allophycocyanin (APC) beads were sorted based on green fluorescence and were treated alongside experimental samples (data not shown). See Supplemental Figure 5 for FACS gating parameters.

#### Pooled cell qRT-PCR

A lysis reaction followed by reverse transcription was performed using the kit Taqman Gene Expression Cells-to-CT Kit (ThermoFisher) with ‘Lysis Solution’ followed by the ‘Stop Solution’ at room temperature, and then a reverse transcription with the ‘RT Buffer’, ‘RT Enzyme Mix’, and lysed RNA at 37°C for an hour. Following reverse transcription, cDNA was pre-amplified by adding 2 uL of cDNA from each sample to 8 uL of preamp master mix [5 uL TaKaRa premix Taq polymerase (Clontech), 2.5 uL 0.2X Taqman pooled probe, 0.5 uL H_2_O] and thermocycled at 95°C for 3 minutes, 55°C for 2 minutes, 72°C for 2 minutes, then 95°C for 15 seconds, 60°C for 2 minutes, 72°C for 2 minutes for 16-20 cycles, and then a final 10°C hold. Amplified cDNA was then diluted 2:100 in RNase free H_2_O. Each qPCR assay contained the following reagents: 0.5 uL 20X Taqman probe, 2.5 uL RNase free H_2_O, 5 uL Gene Expression Master Mix or Fast Advanced Master Mix (ThermoFisher), and 2 uL diluted pre-amplified cDNA. qPCR reactions were performed in triplicate for each Taqman assay of interest on a QuantStudio 3 Real Time PCR Machine (ThermoFisher).

#### qRT-PCR Data Analysis

All three technical replicates were averaged to create one raw Ct values per Taqman assay. As a control, all samples were subject to cell markers of various cell types to confirm specificity: *Rbfox3* for neurons, *Aldh1l1* for astrocytes, *Acta2* for smooth muscle cells, and *Flt1* for endothelial cells (**Supplemental Table 1**). Any assay that did not give a discrete Ct value was given a Ct value of 40 for analysis. Fold change was determined by using the equation 2^(−ΔΔCt)^. *Gapdh* was used as a sample dependent internal control. Fold changes for all runs of each cell type were averaged; the log_2_ of the fold change was calculated, analyzed, and plotted on bar graphs.

#### *In vivo* anesthetized preparation

Animal use was in accordance with guidelines approved by the University of São Paulo Animal Care and Use Committee. A total of 26 adult male C57BL6 (25-28 g) were used for *in vivo* experiments. General anesthesia was induced with 5% isoflurane in 100% O2. A tracheostomy was made, and the isoflurane concentration was reduced to 1.4-1.5% until the end of surgery. The carotid artery was cannulated (polyethylene tubing, 0.6 mm o.d., 0.3 mm i.d., Scientific Commodities) for measurement of arterial pressure (AP). The jugular vein was cannulated for administration of fluids and drugs. Rats were placed supine onto a stereotaxic apparatus (Type 1760; Harvard Apparatus) on a heating pad and core body temperature was maintained at a minimum of 36.5°C via a thermocouple. The trachea was cannulated. Respiratory flow was monitored via a flow head connected to a transducer (GM Instruments) and CO_2_ via a capnograph (CWE, Inc,) connected to the tracheal tube. Paired EMG wire electrodes (AM-System) were inserted into the external intercostal muscle to record respiratory-related activity. After the anterior neck muscles were removed, a basio-occipital craniotomy exposed the ventral medullary surface, and the dura was resected. After bilateral vagotomy, the exposed tissue around the neck and the mylohyoid muscle was covered with mineral oil to prevent drying. Baseline parameters were allowed to stabilize for 30 min prior to recording.

#### In vivo recordings of physiological variables

Mean arterial pressure (MAP), external intercostal muscle activity (IntEMG) and end-expiratory CO_2_ (etCO_2_) were digitized with a micro1401 (Cambridge Electronic Design), stored on a computer, and processed off-line with version 7 of Spike 2 software (Cambridge Electronic Design). Integrated intercostal activity (∫Int_EMG_) was collected after rectifying and smoothing (τ = 0.03) the original signal, which was acquired with a 300-3000 Hz bandpass filter. Noise was subtracted from the recordings prior to performing any calculations of evoked changes in EMG. A direct physiological comparison of the absolute level of EMG activity across nerves is not possible because of non-physiological factors (e.g., muscle electrode contact, size of muscle bundle) and the ambiguity in interpreting how a given increase in voltage in one EMG relates to an increase in voltage in another EMG. Thus, muscle activity was defined according to its baseline physiological state, just prior to their activation. The baseline activity was normalized to 100%, and the percent change was used to compare the magnitude of increases or decreases across muscle from those physiological baselines.

#### In vivo experimental protocol

Each *in vivo* experiment began by testing responses to hypercapnia by adding CO_2_ to the breathing air supplied by artificial ventilation. The addition of CO_2_ was monitored to reach a maximum end-expiratory CO_2_ between 9 and 10%, which corresponds with an estimated arterial pH of 7.2 based on the following equation: pHa = 7.955 − 0.7215 × log10 (EtCO_2_). This maximum end-expiratory CO_2_ was maintained for 5 min and then replaced by 100% O_2_.

To determine whether local regulation of vascular tone in the region of the RTN contributes to the CO_2_/H^+^-dependent drive to breathe, we made injections of saline, PSB1114 (100 µM), AR-C118925 (1 mM) or U46619 (1 µM) while monitoring IntEMG amplitude and frequency. These drugs were diluted with sterile saline (pH 7.4) and applied using single-barrel glass pipettes (tip diameter of 25 µm) connected to a pressure injector (Picospritzer III, Parker Hannifin Corp, Cleveland, OH). For each injection, we delivered a volume of 30 nL over a period of 5s. Injections in the VMS region were placed 1.9 mm lateral from the basilar artery, 0.9 mm rostral from the most rostral hypoglossal nerve rootlet, and at the VMS. The second injection was made 1-2 min later at the same level on the contralateral side. For injections located at the cNTS or ROb, we used the following coordinates: a) cNTS: 0.2-0.3 mm rostral to the *calamus scriptorius*, 0.3 mm lateral to midline, and 0.3 mm below the dorsal surface of the brainstem and ROb: 1.2-1.3 mm caudal to the parietal-occipital suture, in the midline, and 5-5.3 mm below the cerebelar surface.

A cranial optical window was prepared using standard protocols. For the VMS, the anterior neck muscles were removed, a basio-occipital craniotomy exposed the ventral medullary surface, and the dura was resected. Pial vessels in the VMS were located 1.9 mm lateral from the basilar artery and 0.9 mm rostral to the most rostral portion of the hypoglossal nerve rootlet. The surface of the VMS was cleaned with buffer containing (in mmol/L) the following: 135 NaCl, 5.4 KCl, 1 MgCl_2_, 1.8 CaCl_2_, and 5 HEPES, pH 7.3., and a chamber (home-made 1.1-cm-diameter plastic ring was glued with dental acrylic cement attached to a baseplate). The chamber was sealed with a circular glass coverslip (#1943-00005, Bellco). The baseplate was affixed to the Digital Camera (Sony, DCR-DVD3-5) and a light microscope was used for vessel imaging (x40 magnification).

### *In vivo* pial vessel imaging

#### Animal Preparation

Mixed sex adult mice (>6 weeks of age) were briefly anesthetized with isoflurane (1-3%) followed with an IP injection of 300-500mg urethane to ensure a deep anesthetic state. The animal was then fixed to stereotaxic ear bars with the ventral side of the animal facing up. The trachea of the animal was then cannulated with a 18G needle to provide 1.5 mL ventilations at a rate of 150 breaths per minute of 5% CO_2_-21% O_2_ (balance N_2_) via artificial ventilator (Kent Scientific). After cannulation, deep neck muscles were resected, and the cranial-pharyngeal canal and dura mater of the animal was removed. A digital camera was placed over the ventral surface and focused on the pial vasculature. Animals that did not have well perfused pial vessels were not included in experimentation. Animals that were excessively bleeding from the surgical site were sacrificed and not included in any analyses.

#### In vivo Image Analysis

Three linear ROIs were drawn over the pial vasculature we recognize as supplying blood flow to the RTN, similar to the *in vitro* methods described above. Bright field imaging was analyzed using a macro in ImageJ. Briefly, the macro measured the drop and rise of the local maxima and minima, correlating to vessel boundaries. These boundaries were then used to make a linear measurement between the two points which was vessel diameter.

### RTN Viral Injections

Adult mice (>20 grams) were anesthetized with 3% isoflurane. The right cheek of the animal was shaved, and an incision was made to expose the right marginal mandibular branch of the facial nerve. The animals were then placed in a stereotaxic frame and a bipolar stimulating electrode was placed directly adjacent to the nerve. Animals were maintained on 1.5% isoflurane for the remainder of the surgery. An incision was made to expose the skull and two 1.5 mm holes were drilled left and right of the posterior fontanelle, caudal of the lambdoidal suture. The facial nerve was stimulated using a bipolar stimulating electrode to evoke antidromic field potentials within the facial motor nucleus. In this way, the facial nucleus on the right side of the animal was mapped in the X, Y, and Z direction using a quartz recording electrode.

The viral vector was loaded into a 1.2 mm internal diameter borosilicate glass pipette on a Nanoject III system (Drummond Scientific). Virus was injected at least −0.02 mm ventral to the Z coordinates of the facial nucleus, to ensure injection into the RTN. These same coordinates were used for the left side of the animal. In all mice, incisions were closed with nylon sutures and surgical cyanoacrylate adhesive. Mice were placed on a heated pad until consciousness was regained. Meloxicam was administered 24 and 48 h postoperatively.

### Telemetry Transmitter Placement

Adult mice (>30 grams) were anesthetized with an induction dose of 3% isoflurane and placed into a sterile field. HD-X11 transmitters (Data Sciences International; DSI) were placed and fixed intraperitoneally with non-absorbable nylon sutures, followed by ECG lead placement on the left and right pectoralis on top of the rib cage. Finally, a pressure catheter was placed into the left carotid artery and threaded through to the aortic arch for continuous blood pressure measurements. Animals were closed with nylon sutures and placed on a heated pad unit conscious. Mice were continuously watched for the next three days for any post-operative pain complications while meloxicam was administered 24 and 48 h postoperatively. Mice were not used for any experimentation until post-operative day 7.

### Unrestrained whole-body plethysmography

Respiratory activity was measured using a whole-body plethysmograph system (Data Scientific International; DSI), utilizing animal chamber maintained at room temperature and ventilated with air (1.3 L/min) using a small animal bias flow generator. Mice were individually placed into a chamber and allowed 1 hr to acclimate prior to the start of an experiment. Respiratory activity was recorded using Ponemah 5.32 software (DSI) for a period of 15 min in room air followed by exposure to graded increases in CO_2_ from 0% to 7% CO_2_ (balance O_2_). In separate experiments, we characterized the ventilatory response to 10% O_2_ (balance N_2_). Parameters of interests include respiratory frequency (F_R_, breaths per minute), tidal volume (V_T_, measured in mL; normalized to body weight and corrected to account for chamber and animal temperature, humidity, and atmospheric pressure), and minute ventilation (V_E_, mL/min/g). A 20 second period of relative quiescence after 5 min of exposure to each condition was selected for analysis. All experiments were performed between 9 a.m. and 6 p.m. to minimize potential circadian effects.

### Comprehensive Lab Animal Monitoring (CLAMS)

Metabolic monitoring (VCO_2_, VO_2_) was performed using comprehensive lab animal monitoring systems (CLAMS, Columbus Instruments). In short, mice were individually housed on a 12:12 light dark cycle in plastic cages with a running wheel, regular bedding, and regular chow for one week before experimentation. Three days before the metabolic experiment, each animal was placed in the CLAMS housing cage with metered water and waste collection. Mice had two days to acclimate to the metabolic chamber; on the third day, all results were recorded for a continuous 24-hour period. After data collection, all raw results were exported and averaged out per hour. Then, light and dark periods were determined and averaged per animal for statistical analysis. Both sexes were equally represented in the data set.

### Blood Gas Analysis

Arterial blood gasses were collected from adult mice 6 weeks of age and older (>30 grams). A RAPIDLab® 348 blood gas analyzer (Siemens) was used for all blood gas analysis; all calibrations, QC, and use were performed as indicated by the manufacturer. Animals were anesthetized with an induction dose of 3% isoflurane and then quickly switched to 1% isoflurane for the remainder of arterial blood collection. This level of isoflurane minimally affects blood gases (7, 24). The left carotid artery was exposed and quickly cannulated to allow for arterial blood to be collected and analyzed by the blood gas analyzer; no more than 5 seconds was spent between blood collection and analysis on the blood gas analyzer.

### Immunohistochemistry

Adult mice were transcardially perfused with 20 mL of room temperature phosphate buffered saline (PBS, pH 7.4) followed by 20mL of chilled 4% paraformaldehyde (pH 7.4). The brainstem was then removed from the animal and 150 um slices were made using a Zeiss VT100S vibratome. Slices were then incubated in a 0.5% Triton-X/PBS solution for 45 minutes to permeabilize the tissue. The slices remained in a 0.1% Triton-X/ 10% Fetal Bovine Serum (FBS, ThermoFisher)/ PBS solution for a 12-hour primary antibody incubation of anti-mouse alpha smooth muscle actin (Sigma). The tissue was then washed three times in 0.1% Triton-X/10% FBS/PBS solution; the secondary antibody was incubated with the tissue after the third wash for 2 hours (donkey anti-mouse Alexa Fluor 647, ThermoFisher). The tissue was then washed three times in PBS before mounting on precleaned cover slides with Prolong Diamond with DAPI (ThermoFisher). Imaging of brain slices was achieved with a Leica SP8 confocal microscope.

### Statistics

Data are reported as mean ± SE. Power analysis was used to determine sample size, all data sets were tested for normality using Shapiro-Wilk test, and comparisons were made using t-test, one-way or two-way ANOVA (parametric or non-parametric) followed by multiple comparison tests as appropriate. The specific test used for each comparison is reported in the figure legend and all relevant values used for statistical analysis are included in the results section.

## Data Availability

All original data files are available from authors upon request.

## Supplementary Material

### Materials and Methods

**Supplementary Figure 1. CO_2_/H**^**+**^**-induced constriction of RTN arterioles *in vitro* is not dependent on neural activity, prostaglandin EP3 receptors or adenosine signaling. A**, traces of arteriole diameter shows that exposure to 15% CO_2_ alone and in the presence of POM 1 (100 μM; an ectonucleotidase inhibitor), L-798,106 (0.5 μM; a selective EP3 receptor blocker), 8-PT (10 μM; an adenosine receptor antagonist), or TTX (0.5 μM) to block action potentials had similar effects on arteriole diameter. **B**, Summary data show CO_2_-induced change in diameter under control conditions (N=7 vessels) and in the presence of POM 1 (N=7 vessels), L-798,106 (N= 5 vessels), 8-PT (N=8 vessels), and TTX (N=7 vessels) (F_2,19_ = 1.063, p>0.05, one-way ANOVA).

**Supplementary Figure 2. Cardiovascular, metabolic and blood gas parameters in control and smooth muscle P2Y_2_ cKO mice. A**, Traces of blood pressure and ECG activity from control and P2Y_2_ cKO mice. **B**, Summary data (N=4/genotype) shows that control and P2Y_2_ cKO mice have similar mean arterial pressures (MAP) during a 12 hour light cycle (T_3_=0.1152, p > 0.05) and during a 12 hour dark cycle (T_3_=0.1349, p > 0.05). **C**, Summary (N=4/genotype) heart rate (beats/min) during light and dark cycles show similar means for controls and P2Y_2_ cKO (F_1,3_=2.896, p > 0.05). **D-E**, summary (N=4/genotype) blood pressure (**D**) and heart rate (**E**) under room air conditions and during exposure to graded increases in CO_2_ (values were obtained during the last minute of each condition). Control and P2Y_2_ cKO animals show similar blood pressure (F_1,3_=2.051, p > 0.05) and heart rate (F_2,3_=2.896, p > 0.05) responses across all experimental conditions. **F-H**, Oxygen consumption (VO_2_) (**F**), CO_2_ production (VCO_2_) and the respiratory exchange ratio (**H**) were similar between control (N=11 animals) and P2Y_2_ cKO (N=10 animals) mice. **I-K**, under room air conditions control and P2Y_2_ cKO mice (N=7 animals/genotype) showed similar arteriole PO_2_ (T_12_=0.3080, p >0.05), PCO_2_ (T_12_=0.7548, p > 0.05) and pH (T_12_=0.4437, p > 0.05). Blood pressure and heart rate values were compared using a two-way ANOVA with Tukey’s post-hoc multiple comparison test. Baseline metabolic activity and blood gas values were compared using an unpaired two-way t-test.

**Supplementary Figure 3. Respiratory activity of smMHC^Cre/eGFP^::P2Y_2_^f/f^ and control viral injected P2Y_2_ cKO mice.** A-C, We crossed smMHC^Cre/eGFP^ with P2Y^f/f^ as an alternative means of generating smooth muscle specific P2Y_2_ cKO (smMHC^Cre/eGFP^::P2Y_2_^f/f^) mice. Control (smMHC^Cre^ only and P2Y_2_^f/f^ only) and smMHC^Cre/eGFP^::P2Y_2_^f/f^ cKO mice show a similar respiratory frequency (**A**), tidal volume (**B**) and minute ventilation (c) under room air conditions. However, smMHC^Cre/eGFP^::P2Y_2_^f/f^ mice exhibit a blunted ventilatory response to CO_2_. This respiratory phenotype is nearly identical to the respiratory phenotype of αSm22^Cre^::P2Y_2_^f/f^ mice (defined as P2Y_2_ cKO) described in the main text. **D-F**, bilateral RTN injections of control virus (AAV2-Myh11p-eGFP) minimally affected respiratory frequency (**D**), tidal volume (**E**) or minute ventilation (**F**) in αSm22^Cre^::P2Y_2_^f/f^ (P2Y_2_ cKO) mice. *, difference from 0% CO_2_; #, differences between genotypes (two-way ANOVA with Tukey’s multiple comparison test). One symbol = p < 0.05; two symbols = p < 0.01; three symbols = p<0.001; four symbols = p < 0.0001. Linear regressions were compared by two tailed analysis of covariance.

**Supplementary Figure 4. P2Y_2_ cKO mice show a normal ventilatory response to acute hypoxia. A**, Traces of respiratory activity from control and P2Y_2_ cKO mice in room air (21% O_2_) and during exposure to hypoxia (10% O_2_). **B**, Summary data (N=8 animals/genotype) plotted as minute ventilation shows that control and P2Y_2_ cKO exhibit similar respiratory activity under baseline conditions and during hypoxia. ***, different from control at p < 0.001 (two-way RM-ANOVA with Tukey’s multiple comparison test).

**Supplementary Figure 5. Functional characterization of RTN and cortical arterioles in slices from P2Y_2_ cKO mice. A-B**, Diameter traces of individual arterioles and summary data (bottom) show that vessels from the RTN (**A**) and cortex (**B**) in slices from P2Y_2_ cKO mice respond similarly to 15% CO_2_ and t-ACPD (50 μM). As expected, vessels from each region in slices from P2Y_2_ cKO mice failed to respond to bath application of the P2Y_2_ agonist PSB 1114 (200 nM). **C**, RTN arterioles in slices from P2Y_2_ cKO mice two weeks after AAV2-Myh11p-eGFP-2A-mP2ry2 was injected bilaterally into the RTN (P2Y_2_ cKO rescue) constrict in response to 15% CO_2_, t-ACPD and PSB 1114 in a manner similar to vessels from control mice (Fig. 1C). ##, different from baseline (paired two-tailed t-test; p<0.0001).

**Supplementary Figure 6. Cell specificity of AAV2-Myh11p-eGFP-2A-mP2ry2 transduction**. Histological analysis was performed on brainstem sections collected from P2Y_2_ cKO mice two weeks after bilateral RTN injections of AAV2-Myh11p-eGFP-2A-mP2ry2. **Ai**, Example images of virally transduced (eGFP driven by Myh11 promoter) smooth muscle cells counterstained with alpha smooth muscle actin 2 (Acta2; cyan). Overlapping Acta2 and eGFP localization represents positive smooth muscle cell transduction. 4’,6-diamidino-2-phenylindole dihydrochloride (DAPI; white) was used to visualize non-transfected negative cells. **Aii**, Example images of a transduced smooth muscle cell. Some small eGFP green puncta are also observed in a DAPI-positive but Acta2-negative cell (red arrow). Inset, shows eGFP puncta not co-localized with DAPI (yellow arrows). **B**, sagittal view of an arteriole in the RTN with four eGFP labeled smooth muscle cells and some eGFP puncta not co-localized with DAPI (yellow arrows). Scale bars 20 um.

**Supplementary Figure 7. Gating Strategy for FACS sorting of smooth muscle cell and endothelial cell populations from a single cell suspension. A**, Scatter graph gating out debris from the sample. **B**, Side scatter graph to gate for complexity/doublets. **C**, Forward scatter graph to gate for cell size. **D**, Scatter graph for DAPI and GFP (or TdTomato). Cells were gated for positive GFP (or TdTomato) and low DAPI signal.

**Supplementary Table 1. Raw Ct values of housekeeping and cell type specific control genes**. Housekeeping gene *Gapdh* and cell type specific markers including *Rbfox3* (neurons), *Aldh1l1* (astrocytes), *Acta2* (smooth muscle cells), and *Flt1* (endothelial cells) were run in each qRT-PCR panel. Raw Ct values for GAPDH were used for inclusion (cutoff Ct was between 18-22) as well as internal validation.

